# Nucleocapsid mutations in SARS-CoV-2 augment replication and pathogenesis

**DOI:** 10.1101/2021.10.14.464390

**Authors:** Bryan A. Johnson, Yiyang Zhou, Kumari G. Lokugamage, Michelle N. Vu, Nathen Bopp, Patricia A. Crocquet-Valdes, Birte Kalveram, Craig Schindewolf, Yang Liu, Dionna Scharton, Jessica A. Plante, Xuping Xie, Patricia Aguilar, Scott C. Weaver, Pei-Yong Shi, David H. Walker, Andrew L. Routh, Kenneth S. Plante, Vineet D. Menachery

**Affiliations:** Department of Microbiology and Immunology, University of Texas Medical Branch; Galveston, Texas, United States of America; Department of Biochemistry and Molecular Biology, University of Texas Medical Branch; Galveston, Texas, United States of America; Department of Pathology, University of Texas Medical Branch; Galveston, Texas, United States of America; Institute for Human Infection and Immunity, University of Texas Medical Branch; Galveston, Texas, United States of America; World Reference Center of Emerging Viruses and Arboviruses, University of Texas Medical Branch; Galveston, Texas, United States of America; Center for Biodefense and Emerging Infectious Diseases, University of Texas Medical Branch; Galveston, Texas, United States of America; Institute for Drug Discovery, University of Texas Medical Branch; Galveston, Texas, United States of America

## Abstract

While SARS-CoV-2 continues to adapt for human infection and transmission, genetic variation outside of the spike gene remains largely unexplored. This study investigates a highly variable region at residues 203-205 in the SARS-CoV-2 nucleocapsid protein. Recreating a mutation found in the alpha and omicron variants in an early pandemic (WA-1) background, we find that the R203K+G204R mutation is sufficient to enhance replication, fitness, and pathogenesis of SARS-CoV-2. The R203K+G204R mutant corresponds with increased viral RNA and protein both *in vitro* and *in vivo*. Importantly, the R203K+G204R mutation increases nucleocapsid phosphorylation and confers resistance to inhibition of the GSK-3 kinase, providing a molecular basis for increased virus replication. Notably, analogous alanine substitutions at positions 203+204 also increase SARS-CoV-2 replication and augment phosphorylation, suggesting that infection is enhanced through ablation of the ancestral ‘RG’ motif. Overall, these results demonstrate that variant mutations outside spike are key components in SARS-CoV-2’s continued adaptation to human infection.

**Author Summary:** Since its emergence, SARS-CoV-2 has continued to adapt for human infection resulting in the emergence of variants with unique genetic profiles. Most studies of genetic variation have focused on spike, the target of currently available vaccines, leaving the importance of variation elsewhere understudied. Here, we characterize a highly variable motif at residues 203-205 in nucleocapsid. Recreating the prominent nucleocapsid R203K+G204R mutation in an early pandemic background, we show that this mutation is alone sufficient to enhance SARS-CoV-2 replication and pathogenesis. We also link augmentation of SARS-CoV-2 infection by the R203K+G204R mutation to its modulation of nucleocapsid phosphorylation. Finally, we characterize an analogous alanine double substitution at positions 203-204. This mutant was found to mimic R203K+G204R, suggesting augmentation of infection occurs by disrupting the ancestral sequence. Together, our findings illustrate that mutations outside of spike are key components of SARS-CoV-2’s adaptation to human infection.

## Introduction

The emergence of severe acute respiratory syndrome coronavirus 2 (SARS)-CoV-2 is the most significant infectious disease event of the 21st century (1, 2). Since its initial expansion, SARS-CoV-2 has continued to adapt for human infection and transmission, resulting in several variants of concern(3). While most mutations occur within a single lineage, a small number are shared across multiple variants (4). Spike mutations have dominated SARS-CoV-2 variant research, owing to concerns that they enhance replication, augment transmission, or allow escape from immunity (4). However, less attention has been focused on mutations outside spike, despite the existence of other “mutational hotspots” in the genome (4). The SARS-CoV-2 nucleocapsid (N) gene is one hotspot for coding mutations, particularly at amino acid residues 203-205 within its serine rich (SR) domain (5). Three prominent mutations occur in this region including R203K+G204R, a double substitution (KR mt) present in the alpha, gamma, and omicron variants; T205I present in the beta variant; and R203M that occurs in the kappa and delta variants (6, 7). Together, this genetic variation and convergent evolution in residues 203-205 suggests positive selection in this motif of N.

Here, we utilize our reverse genetic system (8, 9) to generate the KR nucleocapsid mutation in the ancestral WA-1 strain of SARS-CoV-2. This change alone was sufficient to increase viral replication in respiratory cells and exhibited enhanced fitness in direct competition studies with wild type (WT) SARS-CoV-2. In the hamster model, the KR mutant (mt) enhanced pathogenesis and outcompeted WT in direct competition. We subsequently found that the KR mt corresponds with increased viral RNA both *in vitro* and *in vivo*. Notably, we observed that the KR mt resulted in augmented nucleocapsid phosphorylation relative to WT SARS-CoV-2; similar increases in N phosphorylation were also seen in the alpha and kappa variants. Importantly, the KR mt was more resistant to GSK-3 kinase inhibition relative to WT SARS-CoV-2, suggesting that the KR mt alters interactions with N targeting kinases. Finally, an analogous alanine double substitution mutant at position 203+204 (AA mt) also increased fitness and altered phosphorylation relative to WT SARS-CoV-2. Together, these results suggest that disruption of the ancestral “RG” motif in nucleocapsid augments infection, fitness, and pathogenesis of SARS-CoV-2.

## Results

### Genetic Analysis of a highly variable motif of SARS-CoV-2 Nucleocapsid

Using SARS-CoV-2 genomic data from the GISAID database(6), we binned each sequence by month of collection and performed an *in silico* search for variation at residues 203-205 within nucleocapsid (**Fig 1A**). Three prominent mutations emerged from this analysis. The first is the R203K+G204R double substitution (KR mt), present in the alpha, gamma, and omicron variants (**Fig 1D**). Historically, the KR mt has been the most abundant mutation in this region, emerging early in the pandemic and reaching 73% of reported sequences in April 2021 (**Fig 1A, S1 Table**). The second prominent mutation, T205I, is present in the beta, eta, and mu lineages (**Fig 1D**). While also emerging early in the pandemic, T205I is a minority variant which peaked at 9% in February 2021 (**Fig 1A, S1 Table**). The third prominent variant mutation is R203M, present in the delta and kappa variants (**Fig 1D**). Interestingly, while the R203M mutation was first detected in March 2020, it persisted as a rare (<1%) variant until April 2021 when it began expanding rapidly, reaching 97% of all reported sequences by November 2021, displacing the KR and T205I mutations (**Fig 1A, S1 Table**). However, with the recent emergence of the omicron variant, the KR mt has regained prominence, displacing R203M as the most common mutation and reaching 93% of all newly reported sequences in January 2022. Together, these data reveal a complex pattern of genetic variation and convergent evolution for residues 203-205 of SARS-CoV-2 N.

**Fig 1.**
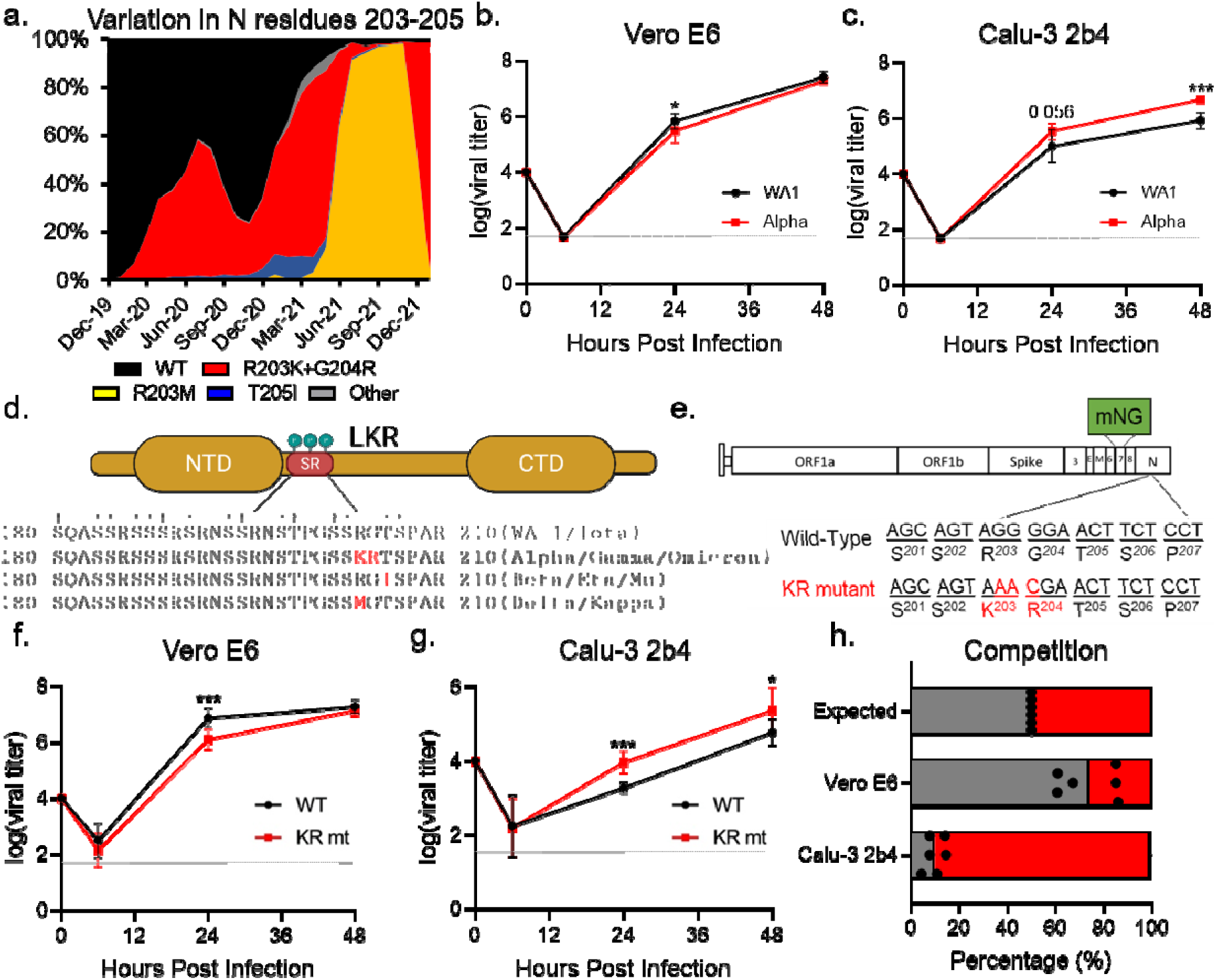
The KR mt enhances SARS-CoV-2 replication. (**A**) Amino acid frequencies for nucleocapsid residues 203-205 in SARS-CoV-2 sequences reported to the GISAID, binned by month of collection and graphed as percent of total sequences reported during that period. (**B-C**) Viral titer from Vero E6 (B) or Calu-3 2b4 cells (C) infected with WA-1 (black) or the alpha variant (red) at an MOI of 0.01 (n≥6). (**D**). Schematic of the SR domain of SARS-CoV-2 nucleocapsid. Variable residues are displayed as red text within the sequence of their corresponding lineages. Phosphorylated residues are indicated by a †. (**E**) Schematic of the SARS-CoV-2 genome, showing the creation of the KR mutation and the replacement of ORF7 with the mNG reporter protein. (**F-G**) Viral titer of Vero E6 (F) or Calu-3 2b4 (G) cells infected with WT (black) or the KR mt (red) at a MOI of 0.01 (n=9). (**H**) Competition assay between WT (gray) and KR mt (red) in Vero E6 or Calu-3 2b4 cells infected at a 1:1 input ratio with an MOI of 0.01 (n=6). Titer data are the mean ± s.d. For competition, individual replicates are graphed as points, while the mean percentage of each virus is displayed as a bar graph. Statistical significance was determined by two-tailed student’s T-test with p≤0.05 (*), p≤0.01 (**), and p≤0.001 (***). Grey dotted lines are equal to LOD.

To determine if mutations in this variable motif have the potential to enhance infection, we evaluated the replication kinetics of SARS-CoV-2 variants. Two cell models were selected for this analysis: Vero E6 (commonly used for propagation and titration of SARS-CoV-2) and Calu-3 2b4 (a respiratory cell line used to study coronavirus and influenza infection) (10, 11). Briefly, Vero E6 or Calu-3 2b4 cells were inoculated at a low multiplicity of infection (MOI) of 0.01 plaque forming units/cell with the early pandemic Washington-1 (WA-1) strain or a SARS-CoV-2 variant and replication kinetics monitored for 48 hours post infection (hpi). In Vero E6 cells, while the alpha and beta variants replicated to equivalent endpoint titers compared to WA-1, both variants had slightly lower titers at 24 hpi (**Fig 1B, S1A Fig**). In contrast, the kappa variant replicated to ∼15-fold lower titer than WA-1 throughout infection (**S1A Fig**), potentially due to processing mutations in SARS-CoV-2 spike shared with the delta variant (12). Interestingly, in Calu-3 2b4 cells, the alpha variant replicated to a 5.6 fold higher endpoint titer compared to WA-1 (**Fig 1C**). In contrast, the beta variant replicated to a lower (2.7 fold) mean endpoint titer compared to WA-1 while the kappa variant showed no significant differences with WA-1 (**S1B Fig**). Together, these data suggest that SARS-CoV-2 variants harboring N mutations may correspond to altered replication kinetics.

### The KR and R203M mts alone are sufficient to increase viral replication

Because the alpha variant exhibited enhanced replication in Calu-3 2b4 cells, we selected the KR mt for further examination. To study the effects of the KR mt in isolation, we utilized a SARS-CoV-2 reverse genetic system to recreate the KR mt in a WA-1 background (**Fig 1E**) (8, 9). In addition, due to gain-of-function concerns, the accessory protein ORF7 was replaced with mNeonGreen (mNG), which reduces but does not eliminate disease in golden Syrian hamsters (**S2 Fig**). After recovery of recombinant virus, we evaluated the KR mt’s effects on SARS-CoV-2 replication. Both Vero E6 and Calu-3 2b4 cells were infected at a low MOI (0.01) with either SARS-CoV-2 WA-1 harboring the mNG reporter (herein referred to as WT) or the KR mt and viral titer monitored for 48 hpi. Like the alpha variant, the KR mt grew to a lower titer at 24 hpi, but had an equivalent endpoint titer in Vero E6 cells (**Fig 1F**). Notably, in Calu-3 2b4 cells, the KR mt had increased viral titer at both 24 and 48 hpi compared to WT SARS-CoV-2 (**Fig 1G**). These data suggest the KR mt in nucleocapsid alone is sufficient to enhance viral replication.

Next, we wanted to examine the ability of other variant mutations in the 203-205 motif of nucleocapsid to enhance SARS-CoV-2 replication. Using our reverse genetic system, we recreated the R203M mutation in a WA-1 mNG background and evaluated its replication in Vero E6 and Calu-3 2b4 cells (**S3 Fig**). Interestingly, like the KR mt, in Vero E6 cells the R203M mutant grew to lower titer at 24 hpi but had a similar endpoint titer compared to WT SARS-CoV-2 (**S3B Fig**). In Calu-3 2b4 cells, the R203M mutant again mimicked the KR mt, growing to a higher titer than WT at 24 and 48 hpi (**S3C Fig**). These data suggest that like the KR mt, the R203M mutation alone is sufficient to enhance SARS-CoV-2 replication.

### The KR mt enhances SARS-CoV-2 fitness during direct competition

We next determined if the KR mt increases SARS-CoV-2 fitness using competition assays, which offer increased sensitivity compared to individual culture experiments (13). WT SARS-CoV-2 and the KR mt were directly competed by infecting Vero E6 and Calu-3 2b4 cells at a 1:1 plaque forming unit ratio. Twenty-four hpi, total cellular RNA was harvested and the ratio of WT to KR mt genomes determined by next generation sequencing (NGS)(14). Consistent with the kinetic data, WT outcompeted the KR mt at a ratio of ∼4:1 in Vero E6 cells (**Fig 1H**). In contrast, the KR mt outcompeted WT at a ratio of ∼10:1 in Calu-3 2b4 cells (**Fig 1H**). These data indicate that the KR mt has a fitness advantage over WT SARS-CoV-2 in Calu-3 2b4, but not Vero E6 cells.

### The KR mt increases pathogenesis and fitness in vivo

We next determined the effects of the KR mt *in vivo* using a golden Syrian hamster model of SARS-CoV-2 infection (15). Three- to four-week-old male hamsters underwent intranasal inoculation with either PBS (mock) or 10^4^ plaque forming units (PFU) of WT SARS-CoV-2 or the KR mt and weight loss was monitored for 10 days post infection (dpi, **Fig 2A**). On days 2 and 4, a cohort of five animals from each group underwent nasal washing, were euthanized, and trachea and lung tissue harvested for measurement of viral loads and histopathologic analysis. On day 10, surviving animals were euthanized and lung tissue harvested for histopathological analysis. Strikingly, animals infected with the KR mt had increased weight loss compared to WT throughout the experiment (**Fig 2B**). Curiously, weight loss changes did not correlate with increased viral loads, as no significant viral titer difference in the lung or trachea was observed between WT and the KR mt (**Fig 2C and D**). Furthermore, the KR mt caused a small, but significant, decrease in titer in nasal washes on day 2, but not day 4 (**Fig 2E**). Contrasting the titer data, histopathologic analysis of lungs revealed that the KR mt had more severe lesions compared to WT SARS-CoV-2 (**S4 Fig**). Compared to mock (**S4A Fig**), both WT and the KR mt had bronchiolitis and interstitial pneumonia; however, larger and more diffuse pulmonary lesions were observed in the KR mt on day 4 (**S4B and C Fig**). In addition, the KR mt had cytopathic alveolar pneumocytes and alveoli containing both mononuclear cells and red blood cells (**S4C Fig**). By day 10, both WT and the KR mt showed signs of recovery, but maintained interstitial pneumonia adjacent to the bronchi absent in mock-infected animals (**S4D–F Fig**). Notably, the KR mt had evidence of cytopathic effect in bronchioles, perivascular edema, and immune infiltration of the endothelium. Together, these data demonstrate that the KR mt increases disease following SARS-CoV-2 infection *in vivo*.

**Fig 2.**
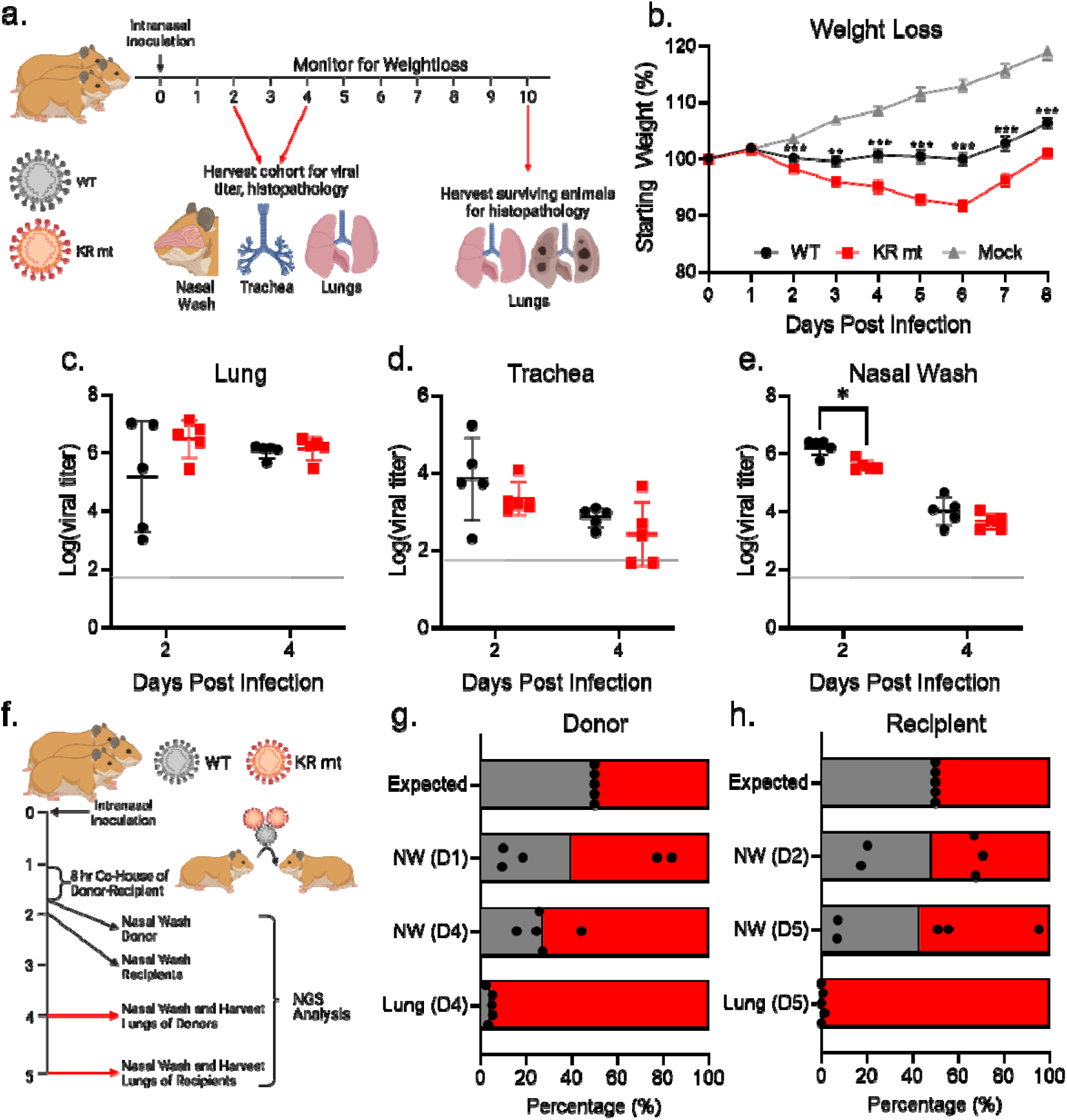
The KR mt enhances SARS-CoV-2 pathogenesis and fitness. (**A**) Schematic of the infection of hamster with SARS-CoV-2. (**B-E**) Three- to four-week-old male hamsters were mock-infected (gray) or inoculated with 10^4^ PFU of WT SARS-CoV-2 (black) or the KR mt (red). Animals were then monitored for weight loss (B). On days 2 and 4 post infection, viral titers in the lung (C), trachea (D), and from nasal washes (E) were determined. (**F**) Schematic of competition/transmission experiment. (**G-H**) Three- to four-week-old male donor hamsters were inoculated with 10^4^ PFU of WT SARS-CoV-2 and the KR mt at a 1:1 ratio. On day 1 of the experiment, donor and recipient hamsters were co-housed for 8 hours, then separated, and the donor hamsters underwent nasal washing. On day 2, recipient hamsters were nasal washed. Hamsters were then monitored and nasal washes and lung tissue harvested on days (donors) and 5 (recipients). The ratio of WT (gray) to KR mt (red) was then determined by NGS of all donor (G) and recipient (H) samples. For weight loss data, mean percent weight loss was graphed ± s.e.m. For titer data, individuals were graphed with means ± s.d. indicated by lines. For competition studies, individual replicates are graphed as points, while bars represent the mean. Statistical significance was between WT and the KR mt was determined by student’s T-Test with p≤0.05 (*), p≤0.01 (**), and p≤0.001 (***). Grey dotted lines are equal to LOD.

We next evaluated the KR mt’s effects on SARS-CoV-2 fitness and transmission *in vivo*. Singly housed three- to four-week-old male donor hamsters were intranasally inoculated with 10^4^ PFU of WT SARS-CoV-2 and the KR mt at a ratio of 1:1 (**Fig 2F**). On day 1 of infection, each donor hamster was co-housed with a recipient for 8 hours to allow transmission. Hamsters were then separated and SARS-CoV-2 present in the nasal cavities of the donors sampled by nasal wash. On day 2, each recipient hamster underwent nasal washing to sample transmitted SARS-CoV-2. Donor and recipient hamsters then underwent nasal washing and harvesting of lung tissue on days 4 and 5, respectively, and the ratio of WT to KR mt in all samples was determined by NGS(14). Neither WT nor the KR mt consistently dominated in the donor washes on day 1; similarly, the day 2 nasal washes from the recipients showed no distinct advantage between KR mt and WT for transmission (**Fig 2G and H**). However, at day 4 and 5 in the donor and recipient, respectively, the KR mt was slightly more predominant in the nasal wash (**Fig 2G and H**). Notably, the KR mt dominated the SARS-CoV-2 population found in lungs of both donor and recipient animals on days 4 and 5. Together, these results suggest that the KR mt outcompetes WT *in vivo* independent of transmission.

### The KR mt increases viral RNA and antigen levels

The CoV N protein has previously been shown to play a role in the transcription of viral RNA (16–19). To evaluate changes in viral RNA levels during infection with the KR mt, we performed RT-qPCR to measure levels of SARS-CoV-2 transcripts following infection of Calu-3 2b4 cells (**S5 Fig**). Compared to WT infected cells, the KR mt had a >32-fold increase in levels of all viral transcripts, demonstrating a broad increase in SARS-CoV-2 RNA levels (**Fig 3A**). This finding is not surprising considering the increased viral titer observed in Calu-3 2b4 cells (**Fig 1G**). We subsequently examined the levels of full-length viral RNA in the lungs of infected hamsters (**Fig 3B**), finding a significant increase at 2- and 4- dpi. In contrast, the virus lung titer at both time points had no significant difference (**Fig 2C**), indicating that the KR mt increases the levels of viral RNA despite not increasing titer. Further extending our analysis, we explored *in vivo* SARS-CoV-2 N antigen staining in the lungs of infected animals (**Fig 3C**). KR mt infected animals showed increased viral antigen staining and substantially larger lesion size compared to WT. Together, these data indicate despite having no effect on lung titer, the KR mt leads to increased viral RNA accumulation and greater virus spread compared to WT SARS-CoV-2.

**Fig 3.**
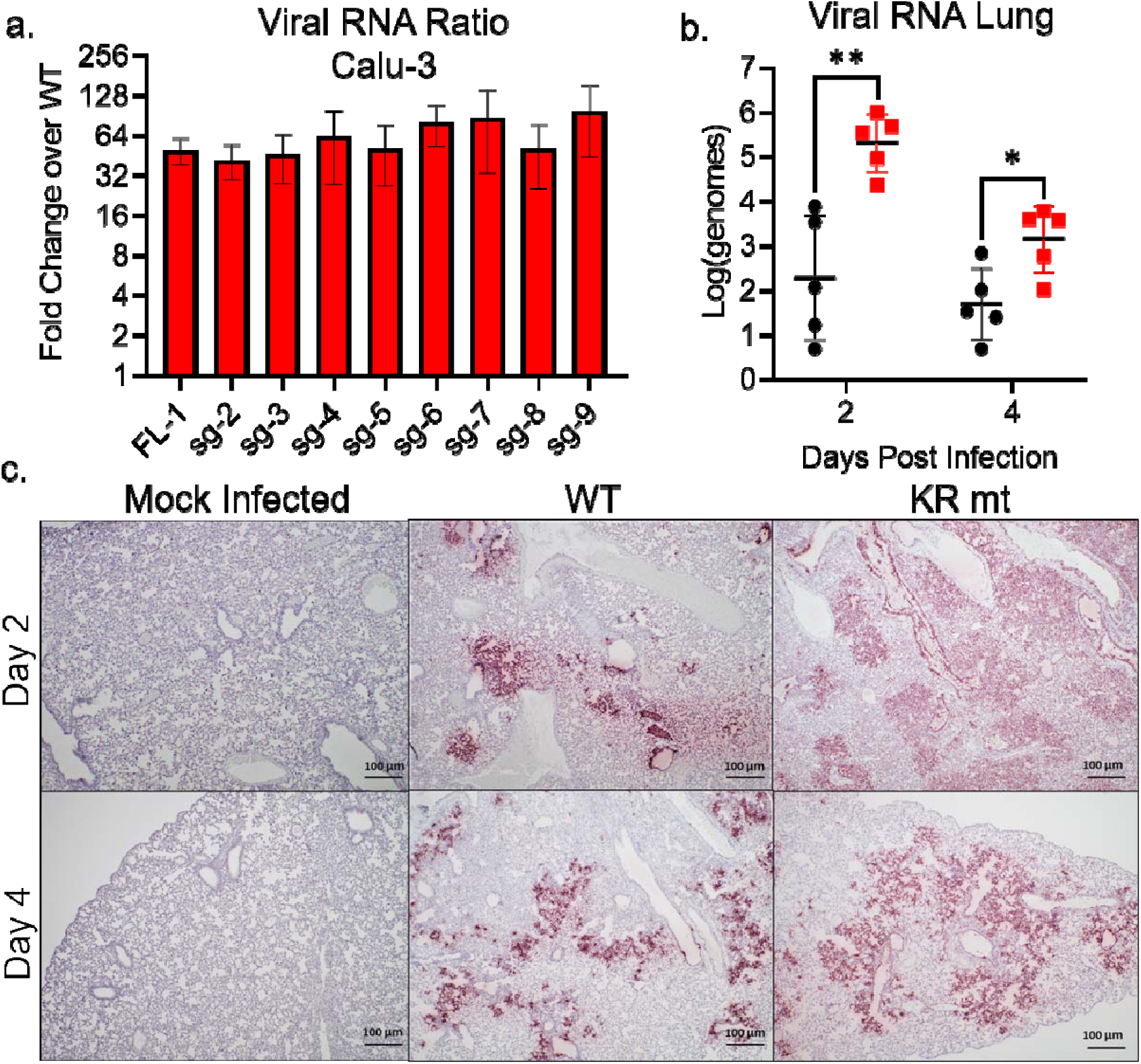
The KR mt increases levels of viral RNA and antigen. (**A**) Full-length and subgenomic transcript levels 2 hours post infection from Calu-3 2b4 cells infected at an MOI of 0.01 with WT SARS-CoV-2 or the KR mt (n=3). Transcript levels were normalized to 18S ribosomal RNA and graphed as the fold change in the KR mt relative to WT (**B-C**) Three- to four-week old male hamsters were inoculated with PBS (mock) or 10^4^ PFU of WT or the KR mt. On days 2 and 4 post infection, lung tissue was harvested. The levels of full-length SARS-CoV-2 RNA in WT and KR mt infected animals (n=5) (B). Representative SARS-CoV-2 antigen staining (anti-Nucleocapsid) of lung tissue from mock, WT, or KR mt infected animals (n=5) (**C**). For *in vitro* transcripts, bars are mean titer ± s.d. For *in vivo* RNA, individual replicates are graphed with means ± s.d. indicated by lines. Significance was determined by student’s T- Test with p≤0.05 (*) and p≤0.01 (**).

### The KR mt increases phosphorylation of SARS-CoV-2 N

Having confirmed a role in viral RNA transcription, we next considered how mutations in residues 203-205 of nucleocapsid’s SR domain might provide an advantage for SARS-CoV-2. Nsp3, the multi-faceted viral protease, interacts with the SR domain to increase viral transcription (18–22). Importantly, this interaction is governed by phosphorylation of the SR domain, which is targeted by the SRPK, GSK-3, and Cdk1 kinases (**Fig 1D**) (5, 22–28) Given the proximity to key priming residues required for GSK-3 mediated phosphorylation (**Fig 4D)** (28), we hypothesized that the KR mt alters nucleocapsid phosphorylation. To overcome the lack of phospho-specific antibodies for nucleocapsid, we used phosphate-affinity SDS-PAGE (PA SDS-PAGE). PA SDS-PAGE utilizes a divalent Zn^2+^ compound (Phos-Tag^TM^) within acrylamide gels that selectively binds to phosphorylated serine, threonine, and tyrosine residues; the bound Zn^2+^ decreases electrophoretic mobility of a protein proportionally with the number of phosphorylated amino acids (29). Importantly, if a protein exhibits multiple phosphorylation states, this will cause a laddering effect, with each phospho-species appearing as a distinct band (**Fig 4A**).

**Fig 4.**
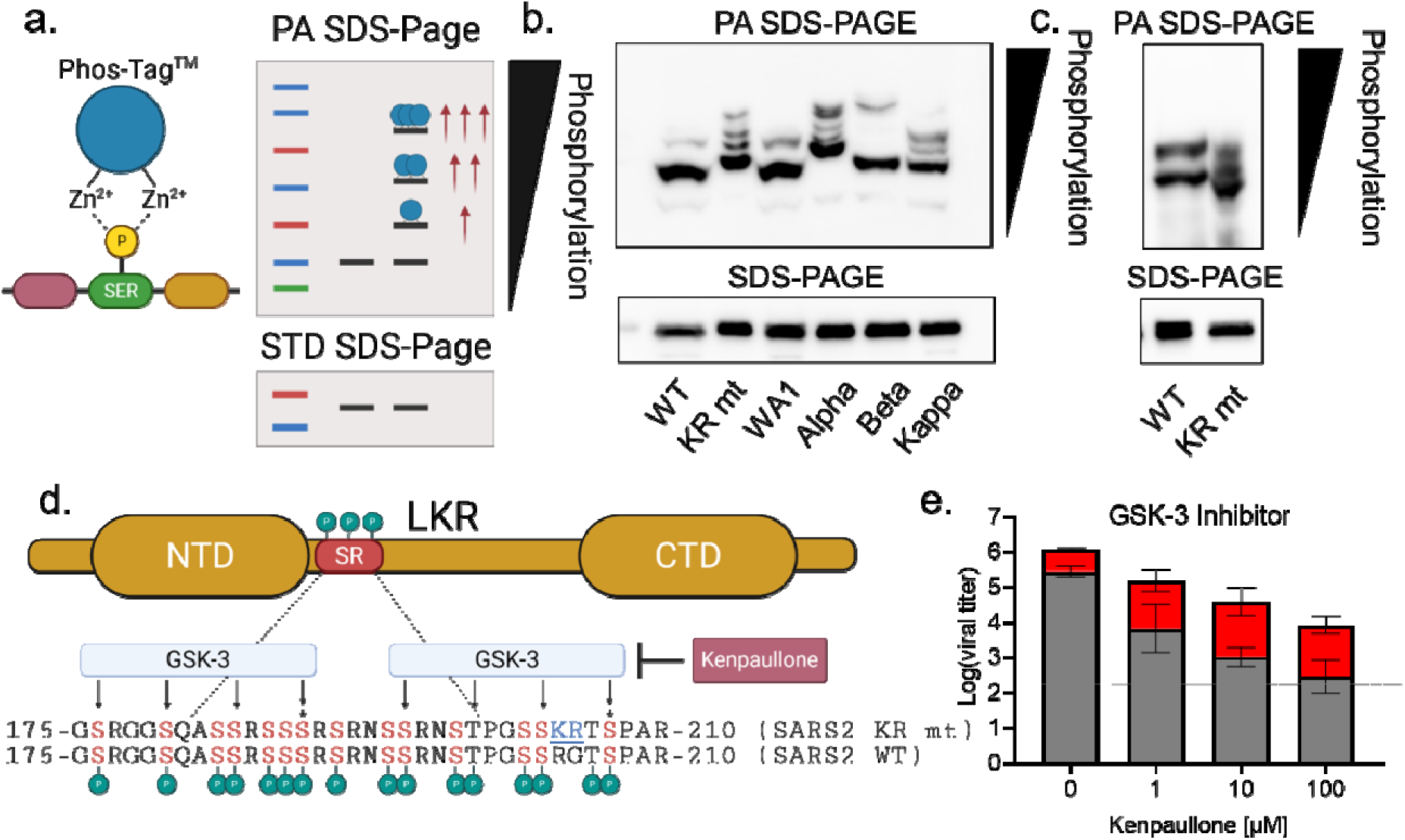
The KR mt increases N phosphorylation to enhance infection. (**A**) Schematic of phosphate-affinity (PA) SDS-PAGE. (**B**) Whole cell lysates from Calu-3 2b4 cells infected with SARS-CoV-2 WA-1-mNG (WT), the KR mt, WA-1, alpha, beta, and kappa variants were analyzed by PA SDS-Page (top) or standard SDS-PAGE (bottom) followed by blotting with an N-specific antibody (n=3). (**C**) Whole cell lysates from Vero E6 cells infected with WT or the KR mt and analyzed by PA-SDS-Page (top) or standard SDS-Page (bottom) followed by blotting with an N-specific antibody (n=3). (**D**) Schematic of phosphorylation by GSK-3 of the SR domain of SARS-CoV-2 N. Residues targeted by GSK-3 are indicated with arrows and priming residues designated by a ‘*’. (**E**) Viral titer 48 hours post infection from Calu-3 2b4 cells infected with WT SARS-CoV-2 (gray) or the KR mt (red) at an MOI of 0.01. Cells were treated with the indicated concentrations of kenpaullone prior to and during infection. Bars are the mean titer ± s.d. (n=4). Significance indicates a change in mean titer difference and was determined by student’s T-Test with p≤0.05 (*) and p≤0.01 (**).

To assess the KR mt’s effects on N-phosphorylation, we infected Calu-3 2b4 cells at a MOI of 0.01 and harvested whole cell lysates 48 hpi. Lysates then underwent PA SDS-PAGE followed by western blotting with an N-specific antibody. When analyzed by PA SDS-PAGE, WT SARS-CoV-2 displayed a two-band pattern consisting of a faint upper and prominent lower band, corresponding to a highly phosphorylated and a less phosphorylated species, respectively (**Fig 4B**, lane 1). In contrast, the KR mt displayed four dark bands of progressively slower mobility, indicating a substantially different phosphorylation pattern (**Fig 4B**, lane 2). Importantly, all four bands migrated more slowly than the prominent WT band, indicating an overall increase in phosphorylation in the KR mt, which corresponds to increased virus replication (**Fig 1G**). We next examined N phosphorylation in Vero E6 cells; cells were infected at a MOI of 0.01 and whole cell lysates taken 24 hpi. In Vero E6 cells, WT SARS-CoV-2 exhibited 2-bands of equal strength, indicating a relative increase in phosphorylation compared to Calu-3 2b4 cells (**Fig 4C**, lane 1). In contrast, the KR mt displayed 3 dark bands with faster mobility relative to WT, indicating a decrease in overall phosphorylation **(Fig 4C**, lane 2). This reduced phosphorylation corresponds with the replication attenuation seen in this cell type (**Fig 1F**). Together, these data indicate that the relative level of SARS-CoV-2 nucleocapsid phosphorylation plays a role in virus replication.

Given the KR mt’s effects, we next determined if SARS-CoV-2 variants had altered N-phosphorylation. Calu-3 2b4 cells were infected at a MOI of 0.01 with WA-1 or the alpha, beta, or kappa variants and whole cell lysates harvested at 48 hpi. When analyzed by PA SDS-PAGE, WA-1 had a two-band pattern similar to WT SARS-CoV-2, while the alpha variant displayed a four-band pattern with slower mobility similar to that of the KR mt (**Fig 4B**, lanes 3-4). Interestingly, the mobilities of the alpha variant bands were decreased compared to the KR mt indicating an even higher level of phosphorylation, potentially due to the alpha variant’s additional nucleocapsid mutations at D3L and S235F(6, 7). While both the beta (T205I) and kappa (R203M) variants also displayed slower electrophoretic mobility compared to WA-1, the beta variant displayed a two-band pattern reminiscent of WA-1 while kappa displayed a laddered pattern similar to the KR mt (**Fig 4B**, lanes 5-6). Together, these data suggest variant mutations at residues 203-205 result in increased N phosphorylation.

### The KR mt does not alter phosphorylation of virion-associated N

While CoV N proteins are hyperphosphorylated intracellularly, they are believed to lack phosphorylation within the mature virion (30, 31). Nevertheless, given the ability of the KR mt to augment phosphorylation, we were curious if it influenced virion-associated SARS-CoV-2 N. To examine this, Calu-3 2b4 cells were infected with WT SARS-CoV-2 or the KR mt. 48 hpi, viral supernatants were taken, clarified, and virions pelleted on a 20% sucrose cushion by ultracentrifugation. Protein recovered from the pellets was then analyzed by PA SDS-PAGE followed by western blotting with an N-specific antibody. Curiously, for both WT and the KR mt, a light upper and dark lower band were detected, indicating some level of N phosphorylation is present in the SARS-CoV-2 virion, albeit at a lower level than intracellular N (**S6 Fig**). However, the KR mt had no effect on the banding pattern, indicating the KR mt does not affect phosphorylation of mature virions.

### The KR mt is more resistant to GSK-3 inhibition

Our results indicate that changes in N phosphorylation correlate with differences in virus replication; thus, we sought to modulate N phosphorylation using kinase inhibitors. Prior work has identified two consensus sites for GSK-3 phosphorylation within the SR domain and inhibition of GSK-3 has been shown to reduce SARS-CoV-2 replication (**Fig 4D**) (28). Importantly, the KR mt is proximal to the priming residue of the C-terminus GSK-3 consensus site, suggesting it may impact GSK-3 mediated N phosphorylation. Therefore, we examined the impact of GSK-3 inhibition on both WT and the KR mt. Using kenpaullone, a GSK-3 inhibitor, we showed a dose dependent inhibitory effect on both WT and KR mt titer at 48 hpi (**Fig 4E**). Importantly, GSK-3 inhibition had a greater impact on WT, significantly increasing the mean titer difference between WT and the KR mt from ∼4-fold to ∼38-fold (**S7 Fig**). This suggests that the KR mt is more resistant to GSK-3 inhibition, and that the change at position 203-204 increases affinity of the KR mt for GSK-3.

### An alanine double substitution mimics the KR mt

Given that variant mutations at residues 203-205 are diverse in sequence, we assessed the importance of the specific R→K and G→R mutations to the KR mt’s enhancement of infection. To do so, we made a R203A+G204A double alanine substitution mutant (AA mt) in the WA-1 mNG background (**S8 Fig**). After recovery of recombinant SARS-CoV-2, we assessed replication in Vero E6 and Calu-3 2b4 cells. In contrast with the KR mt, the AA mt had no significant effect on titers in Vero E6 cells (**Fig 5A**). Nevertheless, the AA mt increased viral titers over WT in Calu-3 2b4 cells throughout infection, mimicking the augmented replication of the KR mt (**Fig 5B**). We next tested the fitness of the AA mt by direct competition with WT SARS-CoV-2. Vero E6 and Calu-3 2b4 cells were infected with WT and the AA mt at a 1:1 ratio, whole cell RNA harvested 24 hpi, and the ratio of WT to AA mt determined by NGS. In Vero E6 cells, the AA mt had a small but consistent fitness advantage over WT (**Fig 5C**). In contrast, the AA mt outcompeted WT with a ∼5:1 ratio during infection of Calu-3 2b4 cells (**Fig 5C**). Overall, the similarities in replication and fitness between the KR and AA mts in Calu-3 2b4 cells suggest that the KR mt enhances infection primarily by ablating the ancestral ‘RG’ motif.

**Fig 5.**
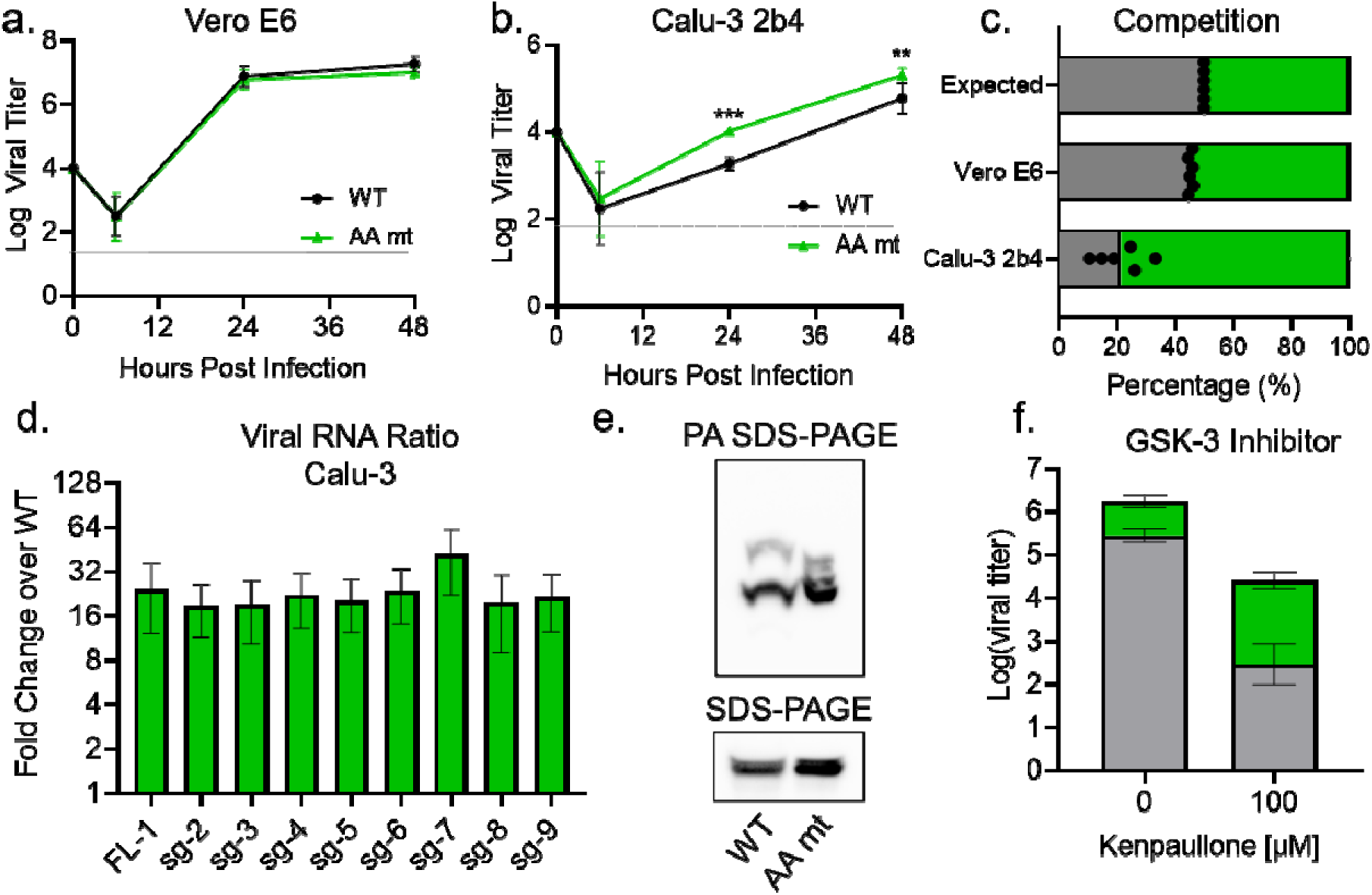
The AA mt mimics the KR mt’s enhancement of SARS-CoV-2 infection. (**A-B**). Viral titer from Vero E6 (A) or Calu-3 2b4 (B) cells infected at an MOI of 0.01 with WT (black) or the AA mt (green) (n=9). (**C**) Competition assay between WT (gray) and the AA mt (green) in Vero E6 and Calu-3 2b4 cells at a 1:1 input ratio and an MOI of 0.01 (n=6). (**D**) Full-length and subgenomic transcript levels 24 hours post infection from Calu-3 2b4 cells infected with WT or the AA mt. Transcripts were normalized to 18S ribosomal RNA and graphed as fold change in the AA mt relative to WT (n=3). (**E**) Whole cell lysates from Calu-3 2b4 cells infecte with WT or the AA mt and analyzed by PA SDS-Page (top) and standard SDS-Page (bottom) followed by blotting with an N-specific antibody (n=3). (**F**) Viral titer 48 hours post infection from Calu-3 2b4 cells infected with WT (gray) or the AA mt (green) at an MOI of 0.01. Cells were treated with the indicated concentrations of kenpaullone prior to and during infection (n=4). Graphs represent mean titer ± s.d. Significance was determined by two-tailed student’s t-test with p≤0.01 (**) and p≤0.001 (***). Grey dotted lines are equal to LOD.

Given that the AA mt mimicked the KR mt’s enhancement of *in vitro* infection, we next determined if they employed the same mechanism. Using RT-qPCR, we examined viral RNA expression in the AA mt (**Fig 5D**). Consistent with the KR mt, at 24 hpi the AA mt increased viral transcript levels compared to WT in Calu-3 2b4 cells, although the magnitude of change in the AA mt was less than the KR mt (16 fold vs 32 fold). We next examined changes in N phosphorylation. Calu-3 2b4 cells were infected at a MOI of 0.01 with WT SARS-CoV-2 or the AA mt and whole cell lysates were harvested 48 hpi. Lysates then underwent PA SDS-PAGE, followed by Western blotting for nucleocapsid. Interestingly, the AA mt exhibited several dark bands with laddered mobility absent in WT, suggesting augmented phosphorylation similar to the KR mt. However, contrasting with the KR mt, in the AA mt the electrophoretic mobility of the lowest band was similar to WT (**Fig 5D**). These data suggest that while the KR and AA mts both alter nucleocapsid phosphorylation, their effects are not identical. Nevertheless, the changes in phosphorylation induced by the KR and AA mt are both sufficient to enhance SARS-CoV-2 replication. Finally, we examined GSK-3 inhibition comparing WT and AA mt (**Fig 5F**). Similar to the KR mt, the AA mt was less sensitive to GSK-3 inhibition than WT, suggesting that disrupting the original RG motif resulted in increased affinity for the kinase. Overall, these data demonstrate that mutations at positions 203-204 disrupt an ancestral motif that interferes with GSK-3 kinase activity.

## Discussion

In this manuscript, we investigate a highly variable motif in SARS-CoV-2 nucleocapsid by characterizing the effects of the R203K+G204R double substitution mutation (KR mt), present in the alpha and omicron variants. Inserting the KR mt in the WA-1 background demonstrated that the KR mt alone is sufficient to increase titer and fitness in respiratory cells. Similarly, the KR mt is sufficient to increase disease in infected hamsters and enhance fitness in the lung during direct competition studies. Importantly, we demonstrate that the KR mt augmented nucleocapsid phosphorylation, which correlated with increased virus replication. We also demonstrate the KR mt is resistant to inhibition of the GSK-3 kinase, mechanistically linking N phosphorylation with phenotypic changes. SARS-CoV-2 variants harboring analogous mutations in the 203-205 motif also exhibit augmented nucleocapsid phosphorylation, suggesting that increasing N phosphorylation is a common mechanism driving variant emergence. Notably, an analogous double alanine substitution (AA mt) also showed increased replication, fitness, altered nucleocapsid phosphorylation, and similar resistance to kinase inhibition. Together, these data suggest that the KR mt and similar mutations enhance SARS-CoV-2 infection by increasing N phosphorylation through disruption of the ancestral “RG” motif.

The KR mt occurs within the serine-rich (SR) domain of nucleocapsid, which has a complex role during SARS-CoV-2 infection. This region of N is hyper-phosphorylated intracellularly, but hypo-phosphorylated within the mature virion (30, 31). Several studies suggest that phosphorylation of the SR domain acts as a biophysical switch, regulating nucleocapsid function through phase separation (27, 32, 33). In the proposed model, unphosphorylated nucleocapsid forms gel-like condensates with viral RNA and the SARS-CoV-2 membrane (M) protein to facilitate genome packaging and virus assembly. In contrast, phosphorylated nucleocapsid forms distinct liquid-like condensates to promote N binding to SARS-CoV-2 nsp3 (27), G3P1 in stress granules (32), and (presumably) other nucleocapsid functions (34). While not tested in the context of infection, this model is consistent with studies demonstrating interactions between N and M(35, 36), G3BP1 and G3BP2 within stress granules(37–39), and phosphorylated N and nsp3 to promote the synthesis of viral RNA(18-22, 30, 40). Within this context, the KR mt may optimize this biomolecular switch for human infection, increasing the amount of phosphorylated nucleocapsid and shifting the overall function of N during infection. Alternatively, the KR mt may impact the interaction between nucleocapsid and host 14-3-3 proteins which bind the SR domain in a phosphorylation dependent manner(41). This interaction is required for cytoplasmic localization of nucleocapsid in SARS-CoV (42). Notably, one 14-3-3 binding site encompasses the 203-205 motif examined in this study,(41) suggesting that the KR mt may enhance infection by altering this interaction.

Overall, in this study we establish that the KR mt enhances SARS-CoV-2 infection relative to WT, increasing viral fitness *in vitro* and *in vivo*, which along with the N501Y mutation(43), likely selected for the emergence of the alpha variant. We also find that the KR mt increases nucleocapsid phosphorylation; coupled with increased resistance to GSK-3 inhibition, these results provide a molecular basis for the KR mt’s effects. Importantly, we show that other variant mutations in this motif also increase nucleocapsid phosphorylation, which may have aided in the emergence of their respective lineages. When taken with the prevalence of mutations in residues 203-205 of the SR domain among circulating variants, these data suggest that mutations increasing nucleocapsid phosphorylation represent a positive selection event for SARS-CoV-2 during adaptation to human infection. Importantly, our work highlights that mutations outside of SARS-CoV-2 spike have significant effects on infection, and must not be overlooked while characterizing mechanisms of variant emergence.

## Materials and Methods

### Construction of Recombinant SARS-CoV-2

The sequence of recombinant wild-type (WT) SARS-CoV-2 is based on the USA-WA1/2020 strain provided by the World Reference Center for Emerging Viruses and Arboviruses (WRCEVA) and originally isolated by the USA Centers for Disease Control and Prevention(10). Recombinant WT SARS-CoV-2 and mutant viruses were created using a cDNA clone and standard cloning techniques as described previously (8, 9). The recombinant SARS-CoV-2 alpha variant (B.1.1.7) was created by introducing twenty-three individual mutations to the WT SARS-CoV-2 infectious clone(12). Construction of WT SARS-CoV-2 and mutant viruses were approved by the University of Texas Medical Branch Biosafety Committee.

### Cell Culture

Vero E6 cells were grown in DMEM (Gibco #11965-092) supplemented with 10% fetal bovine serum (FBS) and 1% antibiotic/antimitotic (Gibco #5240062). Calu-3 2b4 cells were grown in DMEM supplemented with 10% FBS, 1% antibiotic/antimitotic, and 1 mg/ml sodium pyruvate.

### In vitro SARS-CoV-2 infection

Vero E6 and Calu3 2b4 cells were infected with SARS-CoV-2 according to standard protocols described previously (9, 44). Briefly, growth medium was removed, and cells were subsequently infected with SARS-CoV-2 at a multiplicity of infection (MOI) of 0.01 diluted in 100 µl of PBS. Cells were then incubated for 45 minutes at 37°C and 5% CO_2_. After incubation, the inoculum was removed, cells washed three times with PBS, and fresh growth medium returned. For inhibitor experiments, Calu-3 cells were pretreated with 1-100 µM kenpaullone (Sigma-Aldrich #422000) in growth media for 1 hour at 37°C. Cells were then infected with SARS-CoV-2 at an MOI of 0.01 for 45 min at 37°C and 5% CO_2_. Inoculum was then removed, cells were then washed three times with PBS, and fresh growth media with kenpaullone added.

### Focus forming assays

For viral titrations, a focus-forming assay (FFA) was developed by adapting a protocol for a SARS-CoV-2 focus reduction neutralization test (FRNT) published elsewhere (45). Briefly, culture supernatants, nasal washes, or homogenized tissue containing SARS-CoV-2 underwent five 10-fold serial dilutions. 20 µl of the raw sample or each dilution was then used to infect individual wells of 96-well plates containing Vero E6 cells and incubated for 45 minutes at 37°C and 5% CO_2_. After incubation, the inoculum was removed and 100 µl of 0.85% methylcellulose (Sigma# M0512) overlay added to each well and cells incubated for 24 hours at 37°C and 5% CO_2_. After incubation, the overlay was removed, cells washed three times with PBS, and cells fixed with 10% formalin for 30 minutes to inactivate SARS-CoV-2. Cells were then permeabilized by incubating in 0.1% saponin/0.1% bovine serum albumin in PBS for 30 minutes followed by treatment with a nucleocapsid specific antibody (provided by S. Makino). After overnight incubation at 4°C, cells were washed three times with PBS and incubated with a horseradish peroxidase (HRP)-conjugated anti-rabbit secondary antibody (Cell Signaling #7074) for 1 hour at room temperature. The secondary antibody was then removed by washing three times with PBS and foci developed by the application of TruBlue HRP Substrate (Seracare Life Sciences #55100030). Images were then taken with a Cytation 7 cell imaging multi-mode reader (BioTek) and foci counted manually. Prior to inactivation with 10% formalin, all procedures involving the use of infectious SARS-CoV-2 were performed in Biosafety Level 3 (BSL3) or Animal Biosafety Level 3 (ABSL3) facilities at the University of Texas Medical Branch (Galveston, TX).

### In vitro competition assays

Vero E6 or Calu-3 2b4 cells were infected at a 1:1 ratio of WT to mutant SARS-CoV-2, as determined by stock titers. Twenty-four hours post infection, whole cell RNA was harvested by the addition of Triazol reagent (Invitrogen #15596026) and RNA extracted with the RNA Miniprep Plus kit (Zymo Research #R2072) per the manufacturer’s instructions.

### Library preparation, sequencing, and analysis

Extracted RNA samples were prepared for Tiled-ClickSeq libraries(14), with a pool of 396 unique primers. A modified pre-RT annealing protocol was applied in this study. Briefly, a mixture of template RNA, AzDNT/dNTP, and primer pools were heated at 95°C for 2 min; ramped down to 65°C at 1°C/s; incubated at 65°C for 30s; ramped down to 50°C at 0.1°C/s; the rest of RT components were added to libraries while samples were held at 50°C. All subsequent steps followed the previously described method(14), and the final libraries comprising 300-700 bps fragments were pooled and sequenced on a Illumina NextSeq platform with paired-end sequencing.

The raw Illumina data of the Tiled-ClickSeq libraries were processed with previously established bioinformatics pipelines (14). One modification is the introduction of ten wild cards (“N”) covering the KR and AA mutations in the reference genome to allow *bowtie2*(46) to align reads to wild type or variant genomes without bias. PCR duplications were removed using *UMI-tools*(47), and the number of unique reads representing wild type, KR and AA variants were counted thereafter. All raw sequencing data are available in the NCBI Small Read Archive with BioProject ID: PRJNA773399.

### Analysis of publicly available genomic data

SARS-CoV-2 sequences were accessed from the GISAID database on February 14, 2022(6). Sequences were binned by the month during which the sample was collected. The number of R203M, R203K+G204R, T205I, and all other non-wild type options at positions 203-205 were recorded and subtracted from the total number of sequences, with the balance assumed wild type at positions 203-205. Data was then graphed as a percent of total sequences collected in that month.

### Hamster infection studies

For all experiments, golden Syrian hamsters (male, three-to four-week old) were purchased from Envigo. All studies were carried out in accordance with a protocol approved by the UTMB Institutional Animal Care and Use Committee and complied with USDA guidelines in a laboratory accredited by the Association for Assessment and Accreditation of Laboratory Animal Care. Procedures involving infectious SARS-CoV-2 were performed in the Galveston National Laboratory ABSL3 facility.

For pathogenesis studies, animals were housed in groups of five and intranasally inoculated with 10^4^ PFU of WT or KR mt SARS-CoV-2. For competition/transmission studies, animals were intranasally inoculated with 10^4^ PFU of SARS-CoV-2 comprising both WT and the KR mt at a 1:1 ratio based on stock titer. During competition/transmission studies, animals were singly housed throughout the experiment, except during the 8-hour transmission period, when they were housed in groups of two (1 donor and 1 recipient). For all experiments, animals were monitored daily for weight loss and development of clinical disease until the termination of the experiment. Inoculations and other animal manipulations (except weighing) were carried out under anesthesia with isoflurane (Henry Schein Animal Health).

### SDS-PAGE and western blotting

Vero E6 or Calu-3 2b4 cells were infected with SARS-CoV-2 at an MOI of 0.01 and incubated for 24 or 48 hours, respectively. Virus was then inactivated and whole cell lysates taken by the addition of 2× Laemmli buffer (Bio-Rad #161073) + 2-mercaptoethanol (Bio-Rad #1610710) directly to the cell monolayer followed by incubation at 95°C for 15 minutes. For standard SDS-PAGE, 4–20% Mini-PROTEAN TGX Gels (Bio-Rad #4561093) were used for electrophoresis. For phosphate-affinity SDS-PAGE, 7.5% SuperSep^TM^ Phos-Tag^TM^ gels (Wako Chemical #198-17981) were used for electrophoresis. For all gels, protein was transferred to polyvinylidene difluoride (PVDF) membranes and blotted with a SARS-CoV nucleocapsid specific antibody (Novus Biologicals #NB100-56576) followed by horseradish peroxidase (HRP)-conjugated anti-rabbit secondary antibody (Cell Signaling Technology #7074). Images were developed by treating blots for 5 minutes with Clarity Western ECL substrate (Bio-Rad #1705060) followed by imaging on a ChemiDoc MP System (Bio-Rad #12003154). Images were processed with ImageLab 6.0.1 (Bio-Rad #2012931).

### Virion Purification

Calu-3 2b4 cells were infected with WT or the KR mt at an MOI of 0.01. Forty-eight hours post infection; the supernatants were collected and clarified by centrifugation at 1200 rpm. SARS-CoV-2 virions were then pelleted by ultracentrifugation through a 20% sucrose cushion at 26,000 rpm for 3 hours using a Beckman SW32 Ti rotor. Virion pellets were then inactivated in 2X Laemmli buffer (Bio-Rad #161073). N phosphorylation was then analyzed as described in ‘SDS-PAGE and western blotting.’

### Histology

Left lungs (Days 2 and 4) or both lungs (Day 10) were harvested from hamsters and fixed in 10% buffered formalin solution for at least 7 days. After buffer exchange, fixed tissue was then embedded in paraffin, cut into 5 µM sections, and stained with hematoxylin and eosin (H&E) on a SAKURA VIP6 processor by the University of Texas Medical Branch Histology Laboratory. Briefly, fixed tissues were submerged twice in 10% formalin baths, followed by repeated submersion in a series of alcohol baths ranging from 65-100% alcohol. Tissues were then submerged in xylene three times before embedding in paraffin at 60°C. Sections were then cut and mounted on slides, deparaffinized by repeated washing with xylene and alcohol, and then stained with hematoxylin and counterstained with eosin. Alternatively, after mounting slides were deparaffinized and antigen stained in house with a SARS-CoV-2 N specific antibody (Sino Biologicals #40143-R001) at a dilution of 1:30,000 followed by goat anti-rabbit secondary (Vector Labs #BA1000). Signal was developed with ImmPact NovaRED HRP substrate (Vector Labs # SK-4805).

### Real-time Quantitative PCR

For determination of transcript levels in *in vitro* samples, cells were infected as described in ‘In vitro SARS-CoV-2 infection.’ 24 hours post infection, supernatants were discarded and whole cell RNA collected by the addition of Triazol reagent (Invitrogen #15596026). For *in vivo* samples, hamsters were infected as described in ‘hamster infection studies’ and lung lobes placed in RNAlater (Sigma-Aldrich #R0901) and stored at -80°C. Lung lobes were then placed in 1 milliliter Triazol and homogenized with zirconia beads with a MagNA Lyser instrument (Roche Life Science).

RNA from all sources was extracted from Triazol using the Direct-zol RNA Miniprep Plus kit (Zymo Research #R2072) by the manufacturer’s instructions. cDNA was then generated from each RNA sample with the iScript cDNA Synthesis kit (Bio-Rad #1708891). RT-qPCR was performed with Luna Universal qPCR Master Mix (New England Biolabs #M3003) per the manufactures instructions on a CFX Connect instrument (Bio-Rad #1855200). All experiments used an annealing temperature of 51°C. For the analysis of *in vitro* samples, the relative full-length and subgenomic transcript levels between mutants and WT were determined by the delta-delta CT method, with 18S ribosomal RNA serving as an internal control. For *in vivo* samples, the levels of full-length SARS-CoV-2 genomes were quantitated with an 8-point standard curve (1×10^1^ to 1×10^8^ copies per μl).

A common forward primer binding upstream of the transcription regulatory sequence (TRS) leader region was used for all transcripts (ACCAACCAACTTTCGATCTCT). For determination of full-length SARS-CoV-2 genomes, a reverse primer targeting downstream of the TRS leader region was used (CTCGTGTCCTGTCAACGACA). For each sub-genomic (sg) transcript, reverse primers downstream of each TRS body sequence were used: sg2 (TGCAGGGGGTAATTGAGTTCT), sg3 (GCGCGAACAAAATCTGAAGGA), sg4 (AGCAAGAATACCACGAAAGCA), sg5 (ACCGTTGGAATCTGCCATGG), sg6 (GCCAATCCTGTAGCGACTGT), sg7 [mNeonGreen] (TGCCCTCGTATGTTCCAGAAG), sg8 (ACATTCTTGGTGAAATGCAGCT), and sg9 (CCCACTGCGTTCTCCATTCT). The 18S targeting primers used were forward (CCGGTACAGTGAAACTGCGAATG) and reverse (GTTATCCAAGTAGGAGAGGAGCGAG).

## Acknowledgments

We would like to thank Shinji Makino for gifting a nucleocapsid antibody.

## Author contributions

Conceptualization: BAJ, VDM; Formal analysis: BAJ, YZ, JAP, ALR; Funding acquisition: BAJ, YZ, SCW, P-YS, ALR, VDM; Investigation: BAJ, YZ, KGL, MNV, NB, BK, CS, PACV, YL, DS, JAP, XX, DW, KSP; Methodology: BAJ, YZ, PACV, JAP, ALR, SCW, VDM; Project Administration: BAJ, VDM; Supervision: PA, SCW, P-YS, KSP, ALR, VDM, Visualization: BAJ, JAP, DW, Writing – original draft: BAJ, VDM; Writing – review and editing: BAJ, P-YS, PACV, JAP, YL, XX, SCW, DW, ALR, CS, KSP, VDM

## Data Reporting

All raw sequencing data are available in the NCBI Small Read Archive Bioproject ID: PRJNA773399. Raw data available from the corresponding author upon request.

Correspondence and requests for materials should be addressed to V.D.M.

## Supporting Information

**S1 Fig.**
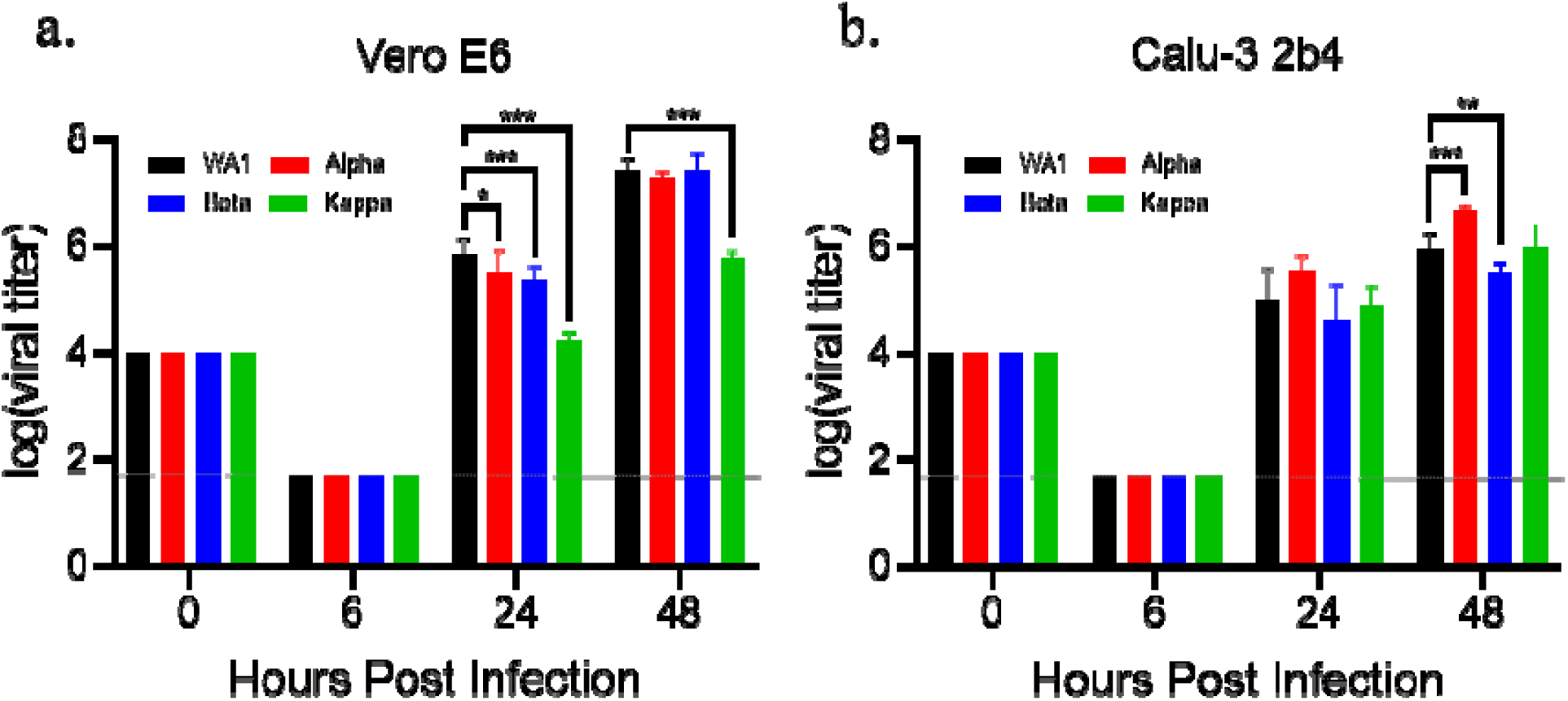
Replication of SARS-CoV-2 variants. Viral titer from Vero E6 (**A**) or Calu-3 2b4 cells (**B**) inoculated with SARS-CoV-2 WA-1 (black) or the alpha (red), beta (blue) or kappa (green) variants at a MOI of 0.01. Graphed dat represent the mean ± s.d. Statistical significance was determined by two-tailed student’s T-test with p≤0.05 (*), p≤0.01 (**), and p≤0.001 (***). Grey dotted lines are equal to LOD.

**S2 Fig.**
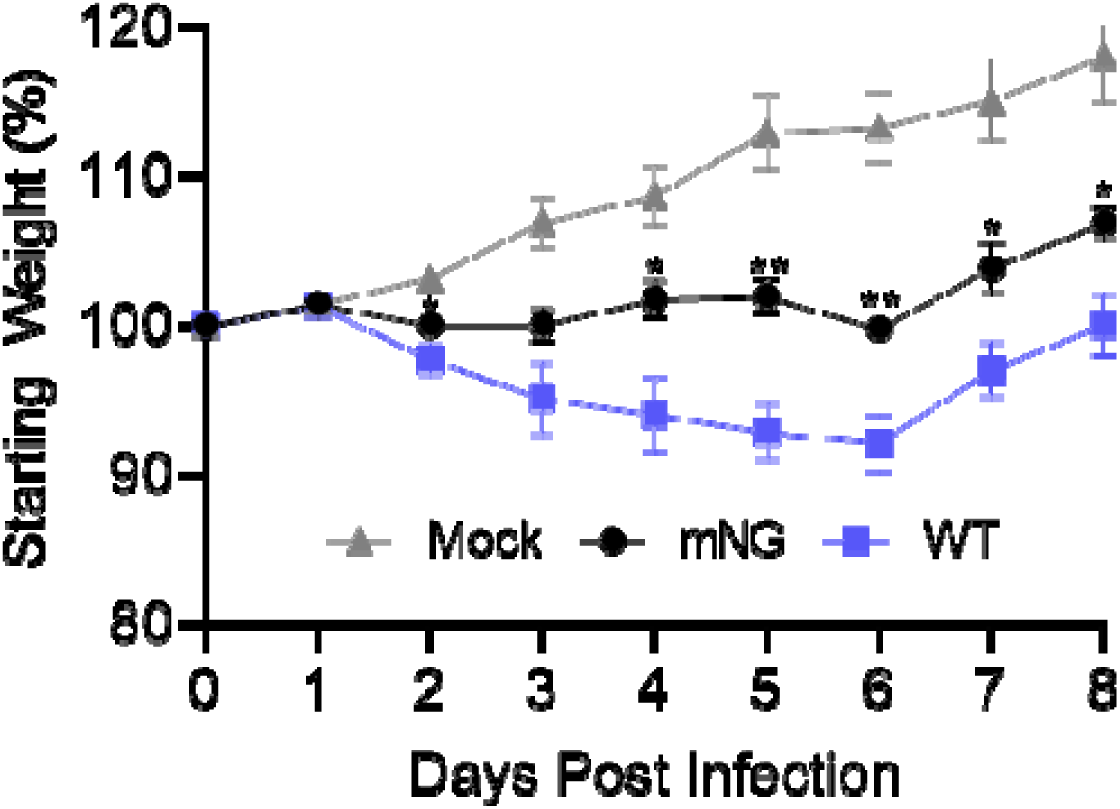
*In vivo a*ttenuation of the SARS-CoV-2 mNeonGreen reporter virus. Three- to four-week-old Golde Syrian hamsters were intranasally inoculated with PBS alone (gray) or 10^4^ PFU of WA-1 SARS-CoV-2 (blue) or mNG SARS-CoV-2 (black). Graphed data represent the mean weight loss ± s.e.m (n≥5). Statistical significance between WT and mNG determined by two-tailed students T-test with p≤0.05 (*) and p≤0.01 (**).

**S3 Fig.**
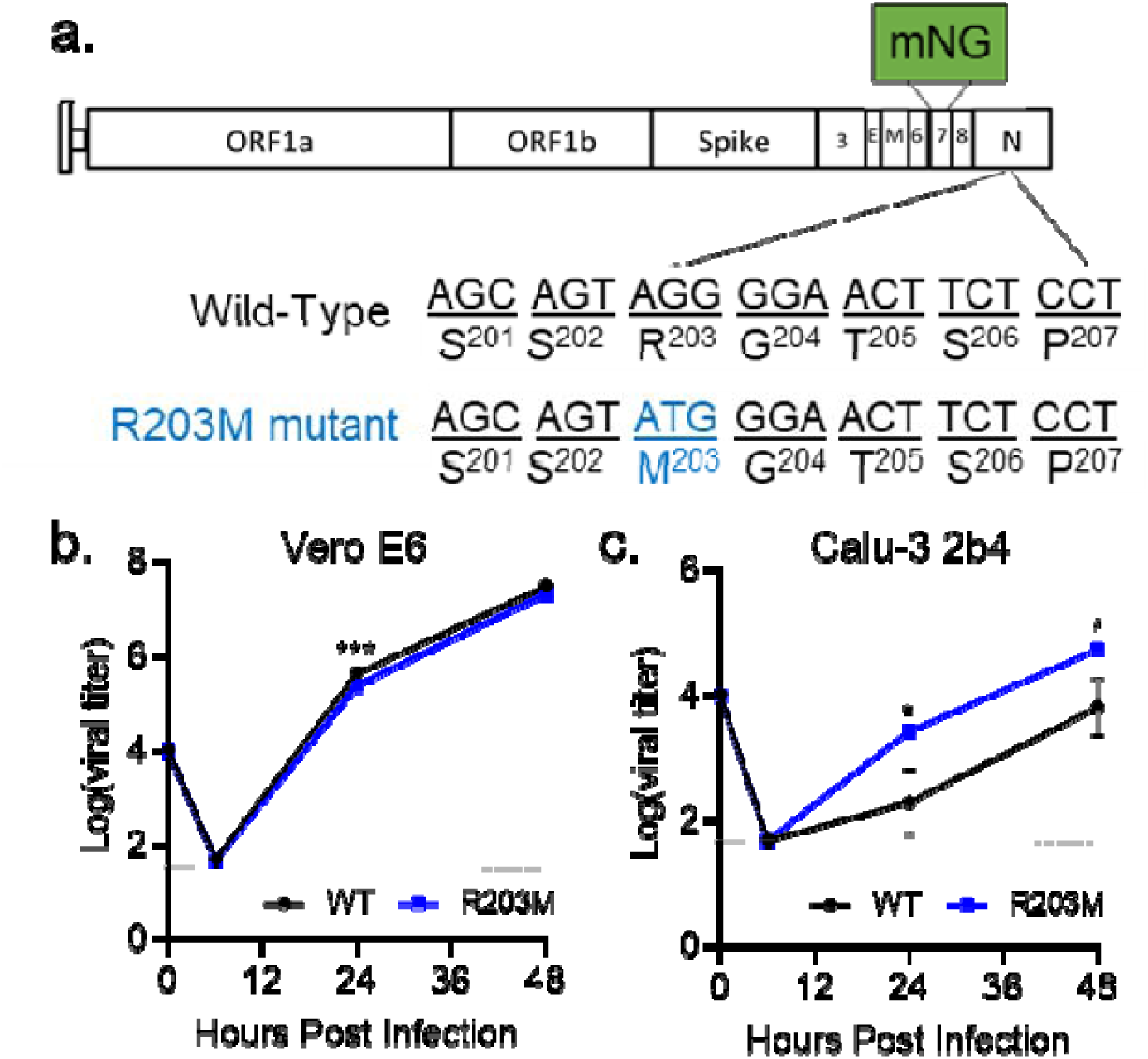
The R203M mutation enhances SARS-CoV-2 replication. (**A**) Schematic of the SARS-CoV-2 genome, showing the creation of the R203M mutation and the replacement of ORF7 with the mNG reporter protein. (**B-C**) Viral titers from Vero E6 (B) or Calu-3 2b4 (C) infected with WT or R203M SARS-CoV-2 at an MOI of 0.01. Graphed data represent mean ± s.d. (n=3). Statistical significance was determined by two-tailed student’s T-test with p≤0.05 (*) and p≤0.001 (***). Grey dotted lines are equal to LOD.

**S4 Fig.**
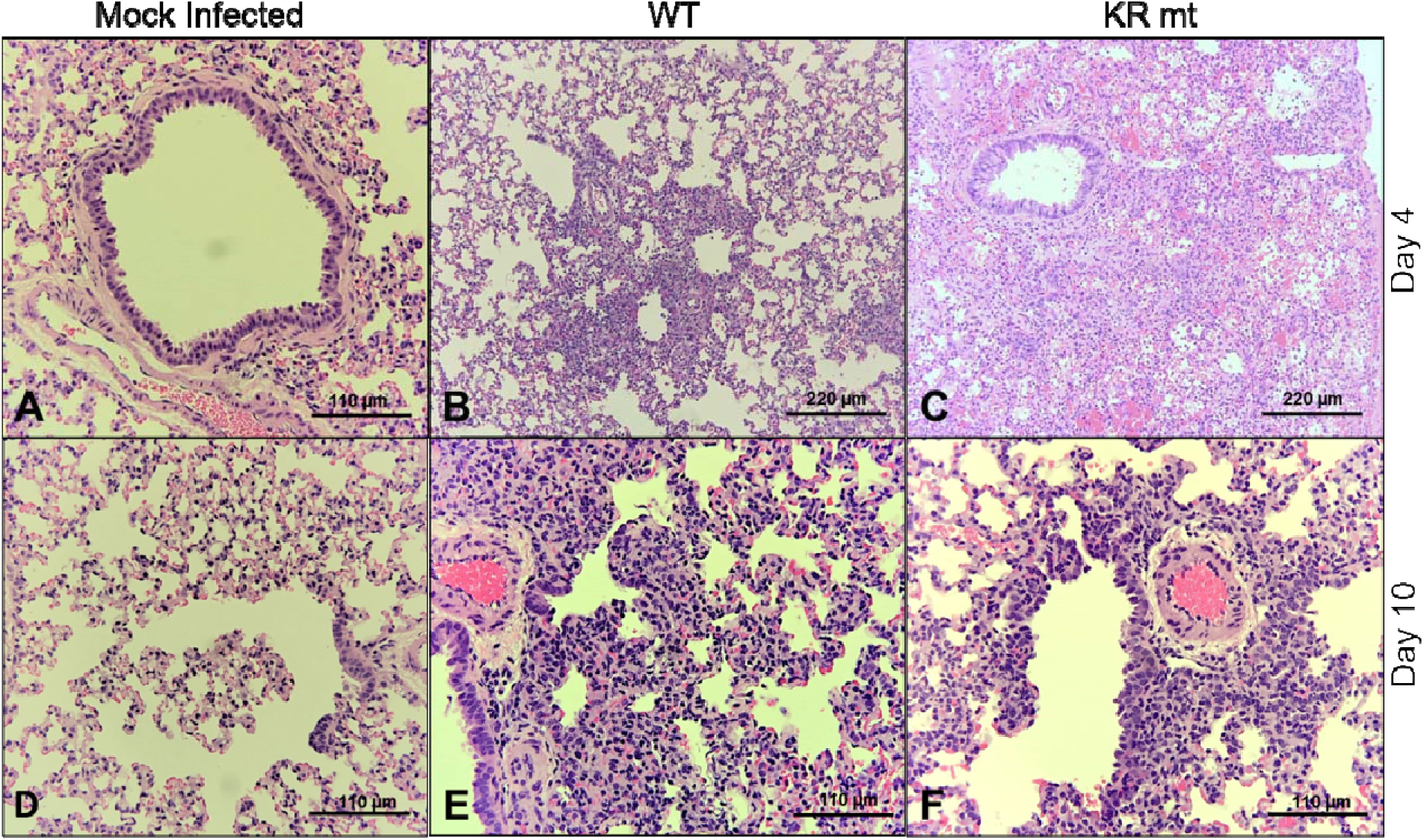
Lung histopathology in hamsters infected with WT and KR mt SARS-CoV-2. Lung tissue was harvested, fixed, and 5 µm sections cut from mock, WT SARS-CoV-2, or KR mt-infected hamsters and stained with hematoxyli and eosin. (**A**) Normal bronchus, pulmonary artery, and alveoli in mock infection on day 4 (20X). (**B**) Bronchiolitis, peribronchiolitis, interstitial pneumonia, and edema surrounding branch of the pulmonary artery at day 4 in hamsters infected with WT virus (10X). (**C**) Severe bronchiolar cytopathic effect, interstitial pneumonia, cytopathic alveolar pneumocytes, alveoli containing mononuclear cells and red blood cells at day 4 in hamsters infected with KR mt. This lesion extended over numerous fields (10X). (**D**) Normal respiratory bronchiole, alveolar ducts, and alveolar sacs in mock infection on day 10 (20X). (**E**) Interstitial pneumonia adjacent to a bronchus at day 10 in hamsters infected with WT (20X). (**F**) Bronchiolar epithelial cytopathic effect, peribronchiolitis, focal interstitial pneumonia, branch of pulmonary artery with surrounding edema and mononuclear cell infiltration of endothelium at day 10 in a hamster infected with the KR mt (20X). Shown are representative images typical of data gathered from 5 animals from each group.

**S5 Fig.**
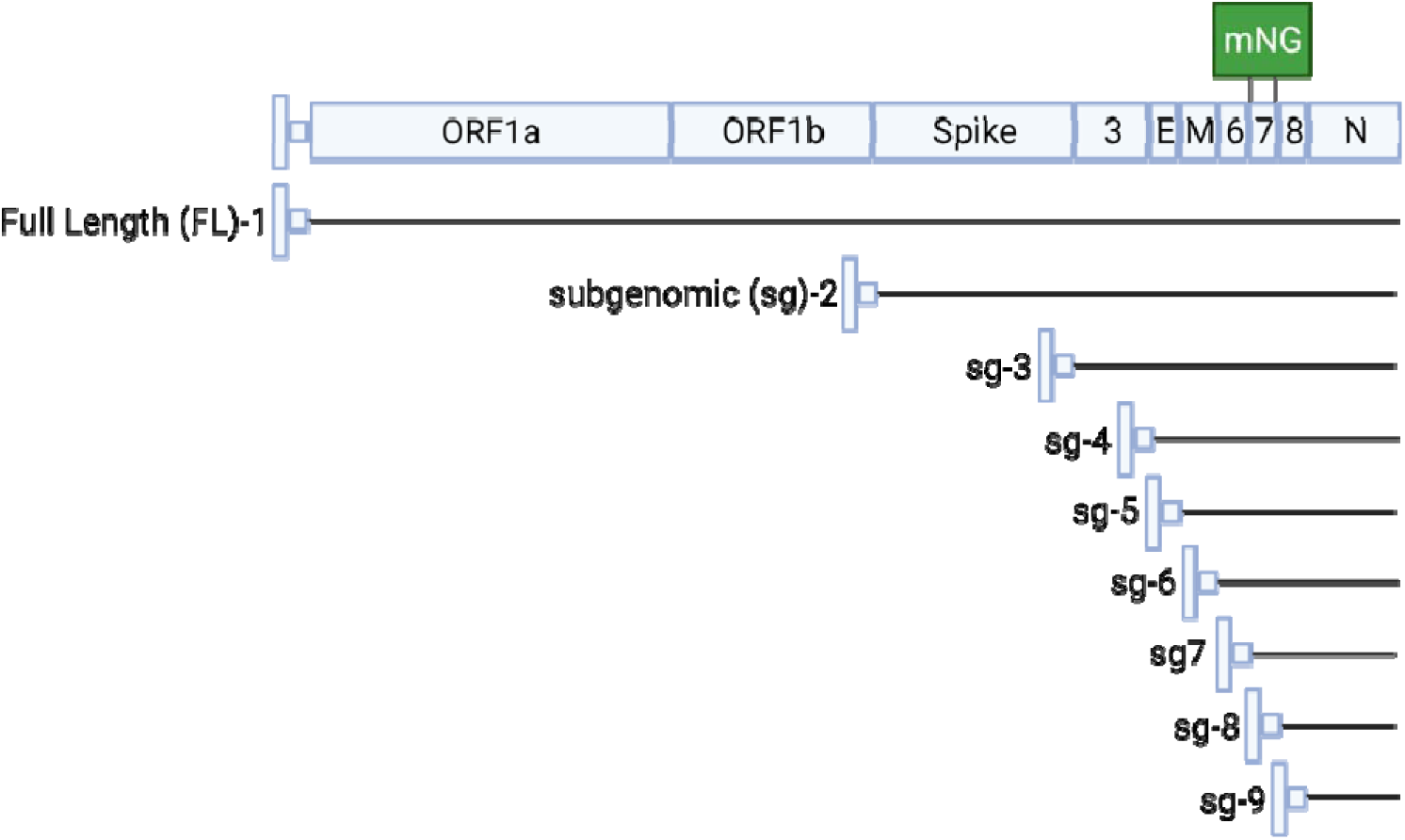
Schematic of SARS-CoV-2 RNAs. Illustration of full length (FL) and subgenomic (sg) RNAs produced during SARS-CoV-2 infection.

**S6 Fig.**
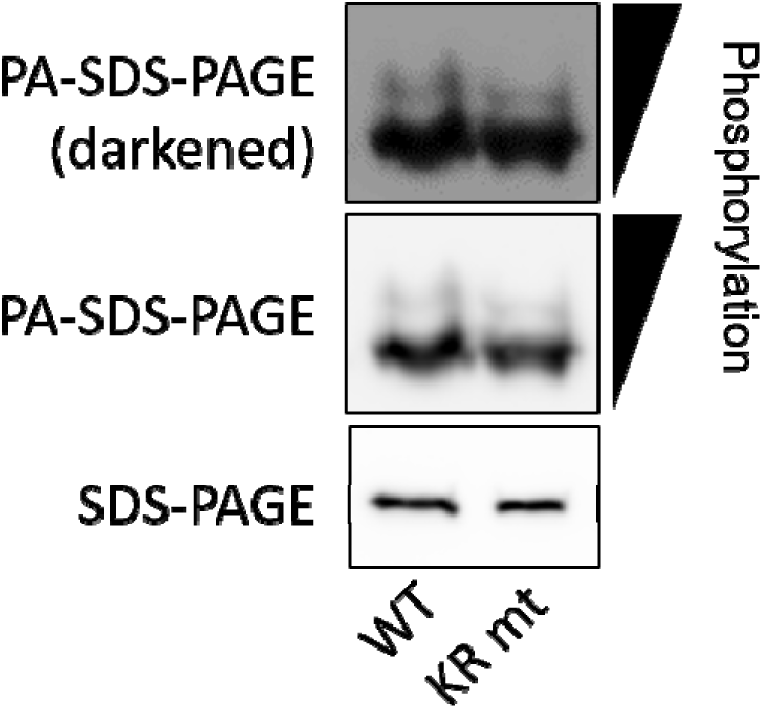
The KR mt has no effect on phosphorylation in virions. Calu-3 2b4 cells were infected at an MOI of 0.0 with WT or KR mt SARS-CoV-2. Forty-eight hours post infection, viral supernatants were taken. Virions were the purified from supernatants by ultracentrifugation on a 20% sucrose cushion, inactivated, and N levels analyzed by both phospho-affinity and standard SDS-Page. Results are representative of two independent experiments.

**S7 Fig.**
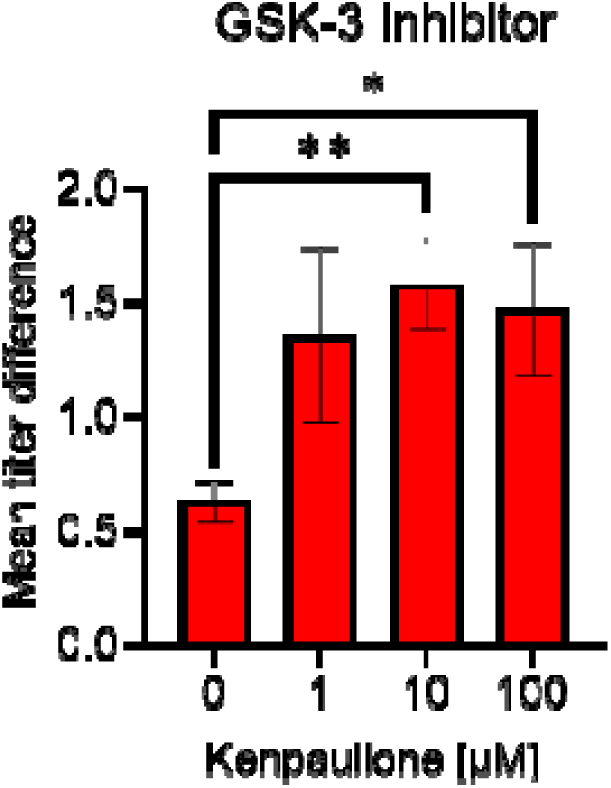
Mean titer differences between WT and KR mutant during GSK-3 inhibition. Mean difference in viral titer between WT SARS-CoV-2 and the KR mutant when treated with kenpaullone at the indicated concentrations (n=4). Error bars are ± s.e.m. Significance by student T-test with p≤0.05 (*) and p≤0.01 (**).

**S8 Fig.**
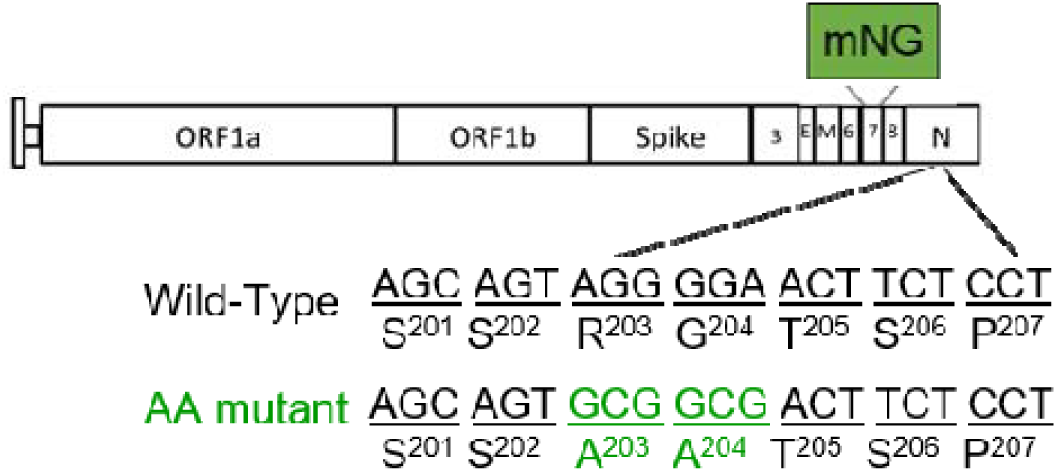
Schematic representation of the AA mt. Schematic shows the creation of the AA mutation within the SARS-CoV-2 genome and the replacement of ORF7 with the mNeonGreen reporter.

**S1 Table.**
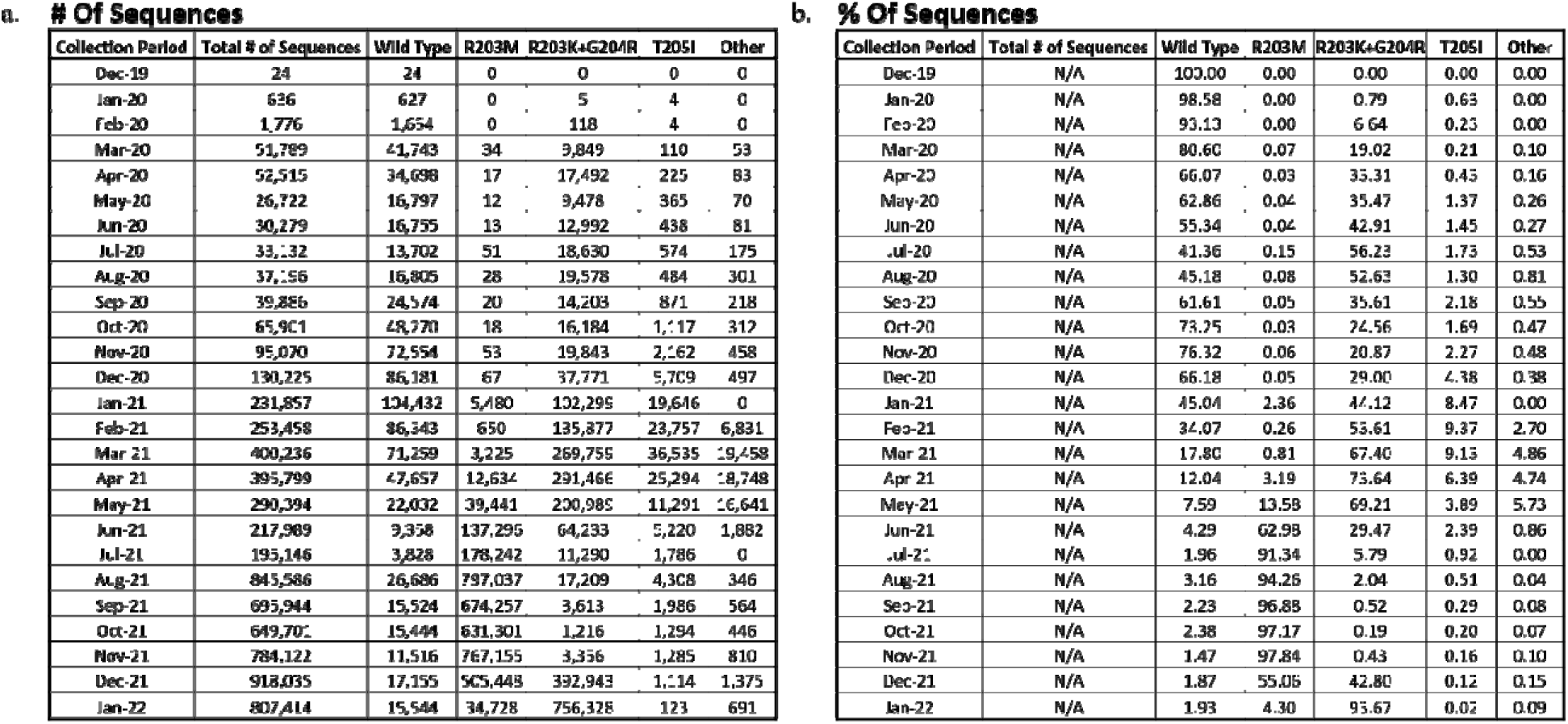
Frequency of mutations in residues 203-205 in SARS-CoV-2 nucleocapsid. (**A-B**) Frequency of WT, R203M, R203K+G204R, T205I, or all other genotypes binned by month of collection, represented as the raw totals (A) or as a percentage of total sequences in a given month (B)..

## Notes

### Competing Interest Statement

XX, P-YS, and VDM have filed a patent on the reverse genetic system and reporter SARS-CoV-2. Other authors declare no competing interests.

### Summary of Updates

Additional mechanistic data has been added.

## References

1. International Monetary Fund Research Dept. World Economic Outlook, April 2020 : The Great Lockdown. Paper. International Monetary Fund, Dept. R; 2020 April 14, 2020.

2. Word Health Organization. Coronavirus disease (COVID-19) pandemic. 2021;2021(29 September).

3. Word Health Organization. Tracking SARS-CoV-2 variants. 2022;2022(17 February).

4. Plante JA, Mitchell BM, Plante KS, Debbink K, Weaver SC, Menachery VD. The variant gambit: COVID-19’s next move. Cell Host Microbe. 2021.

5. Ye Q, West AMV, Silletti S, Corbett KD. Architecture and self-assembly of the SARS-CoV-2 nucleocapsid protein. Protein Sci. 2020;29(9):1890–901.

6. Elbe S, Buckland-Merrett G. Data, disease and diplomacy: GISAID’s innovative contribution to global health. Glob Chall. 2017;1(1):33–46.

7. Mullen JL, Tsuen G, Latif AA, Alkuzweny M, Cano M, Haag E, et al. outbreak.info 2020 [cited 2021 1 October 2021]. Available from: https://outbreak.info/

8. Xie X, Lokugamage KG, Zhang X, Vu MN, Muruato AE, Menachery VD, et al. Engineering SARS-CoV-2 using a reverse genetic system. Nature Protocols. 2021;16(3):1761–84.

9. Xie X, Muruato A, Lokugamage KG, Narayanan K, Zhang X, Zou J, et al. An Infectious cDNA Clone of SARS-CoV-2. Cell Host & Microbe. 2020;27(5):841–8.e3.

10. Harcourt J, Tamin A, Lu X, Kamili S, Sakthivel SK, Murray J, et al. Severe Acute Respiratory Syndrome Coronavirus 2 from Patient with Coronavirus Disease, United States. Emerg Infect Dis. 2020;26(6):1266–73.

11. Menachery VD, Eisfeld AJ, Schäfer A, Josset L, Sims AC, Proll S, et al. Pathogenic influenza viruses and coronaviruses utilize similar and contrasting approaches to control interferon-stimulated gene responses. mBio. 2014;5(3):e01174–14.

12. Liu Y, Liu J, Johnson BA, Xia H, Ku Z, Schindewolf C, et al. Delta spike P681R mutation enhances SARS-CoV-2 fitness over Alpha variant. bioRxiv. 2021:2021.08.12.456173.

13. Plante JA, Liu Y, Liu J, Xia H, Johnson BA, Lokugamage KG, et al. Spike mutation D614G alters SARS-CoV-2 fitness. Nature. 2021;592(7852):116–21.

14. Jaworski E, Langsjoen RM, Michell B, Judy B, Newman P, Plante JA, et al. Tiled-ClickSeq for targeted sequencing of complete coronavirus genomes with simultaneous capture of RNA recombination and minority variants. Elife. 2021.

15. Imai M, Iwatsuki-Horimoto K, Hatta M, Loeber S, Halfmann PJ, Nakajima N, et al. Syrian hamsters as a small animal model for SARS-CoV-2 infection and countermeasure development. Proceedings of the National Academy of Sciences. 2020;117(28):16587.

16. Baric RS, Nelson GW, Fleming JO, Deans RJ, Keck JG, Casteel N, et al. Interactions between coronavirus nucleocapsid protein and viral RNAs: implications for viral transcription. J Virol. 1988;62(11):4280–7.

17. Stohlman SA, Baric RS, Nelson GN, Soe LH, Welter LM, Deans RJ. Specific interaction between coronavirus leader RNA and nucleocapsid protein. J Virol. 1988;62(11):4288–95.

18. Verheije MH, Hagemeijer MC, Ulasli M, Reggiori F, Rottier PJ, Masters PS, et al. The coronavirus nucleocapsid protein is dynamically associated with the replication-transcription complexes. J Virol. 2010;84(21):11575–9.

19. Zuniga S, Cruz JL, Sola I, Mateos-Gomez PA, Palacio L, Enjuanes L. Coronavirus nucleocapsid protein facilitates template switching and is required for efficient transcription. J Virol. 2010;84(4):2169–75.

20. Hurst KR, Koetzner CA, Masters PS. Characterization of a critical interaction between the coronavirus nucleocapsid protein and nonstructural protein 3 of the viral replicase-transcriptase complex. J Virol. 2013;87(16):9159–72.

21. Hurst KR, Ye R, Goebel SJ, Jayaraman P, Masters PS. An interaction between the nucleocapsid protein and a component of the replicase-transcriptase complex is crucial for the infectivity of coronavirus genomic RNA. J Virol. 2010;84(19):10276–88.

22. Koetzner CA, Hurst-Hess KR, Kuo L, Masters PS. Analysis of a crucial interaction between the coronavirus nucleocapsid protein and the major membrane-bound subunit of the viral replicase-transcriptase complex. Virology. 2022;567:1–14.

23. Bouhaddou M, Memon D, Meyer B, White KM, Rezelj VV, Correa Marrero M, et al. The Global Phosphorylation Landscape of SARS-CoV-2 Infection. Cell. 2020;182(3):685–712 e19.

24. Davidson AD, Williamson MK, Lewis S, Shoemark D, Carroll MW, Heesom KJ, et al. Characterisation of the transcriptome and proteome of SARS-CoV-2 reveals a cell passage induced in-frame deletion of the furin-like cleavage site from the spike glycoprotein. Genome Medicine. 2020;12(1):68.

25. Klann K, Bojkova D, Tascher G, Ciesek S, Münch C, Cinatl J. Growth Factor Receptor Signaling Inhibition Prevents SARS-CoV-2 Replication. Mol Cell. 2020;80(1):164–74.e4.

26. Yaron TM, Heaton BE, Levy TM, Johnson JL, Jordan TX, Cohen BM, et al. The FDA-approved drug Alectinib compromises SARS-CoV-2 nucleocapsid phosphorylation and inhibits viral infection in vitro. bioRxiv. 2020:2020.08.14.251207.

27. Carlson CR, Asfaha JB, Ghent CM, Howard CJ, Hartooni N, Safari M, et al. Phosphoregulation of Phase Separation by the SARS-CoV-2 N Protein Suggests a Biophysical Basis for its Dual Functions. Mol Cell. 2020;80(6):1092–103.e4.

28. Liu X, Verma A, Garcia G, Jr., Ramage H, Lucas A, Myers RL, et al. Targeting the coronavirus nucleocapsid protein through GSK-3 inhibition. Proc Natl Acad Sci U S A. 2021;118(42).

29. Kinoshita E, Kinoshita-Kikuta E, Koike T. The Cutting Edge of Affinity Electrophoresis Technology. Proteomes. 2015;3(1):42–55.

30. Wu CH, Chen PJ, Yeh SH. Nucleocapsid phosphorylation and RNA helicase DDX1 recruitment enables coronavirus transition from discontinuous to continuous transcription. Cell Host Microbe. 2014;16(4):462–72.

31. Wu CH, Yeh SH, Tsay YG, Shieh YH, Kao CL, Chen YS, et al. Glycogen synthase kinase-3 regulates the phosphorylation of severe acute respiratory syndrome coronavirus nucleocapsid protein and viral replication. J Biol Chem. 2009;284(8):5229–39.

32. Lu S, Ye Q, Singh D, Cao Y, Diedrich JK, Yates JR, 3rd, et al. The SARS-CoV-2 nucleocapsid phosphoprotein forms mutually exclusive condensates with RNA and the membrane-associated M protein. Nat Commun. 2021;12(1):502.

33. Cubuk J, Alston JJ, Incicco JJ, Singh S, Stuchell-Brereton MD, Ward MD, et al. The SARS-CoV-2 nucleocapsid protein is dynamic, disordered, and phase separates with RNA. Nat Commun. 2021;12(1):1936.

34. McBride R, van Zyl M, Fielding BC. The coronavirus nucleocapsid is a multifunctional protein. Viruses. 2014;6(8):2991–3018.

35. He R, Leeson A, Ballantine M, Andonov A, Baker L, Dobie F, et al. Characterization of protein-protein interactions between the nucleocapsid protein and membrane protein of the SARS coronavirus. Virus Res. 2004;105(2):121–5.

36. Hurst KR, Kuo L, Koetzner CA, Ye R, Hsue B, Masters PS. A major determinant for membrane protein interaction localizes to the carboxy-terminal domain of the mouse coronavirus nucleocapsid protein. J Virol. 2005;79(21):13285–97.

37. Gordon DE, Jang GM, Bouhaddou M, Xu J, Obernier K, White KM, et al. A SARS-CoV-2 protein interaction map reveals targets for drug repurposing. Nature. 2020;583(7816):459–68.

38. Li J, Guo M, Tian X, Wang X, Yang X, Wu P, et al. Virus-Host Interactome and Proteomic Survey Reveal Potential Virulence Factors Influencing SARS-CoV-2 Pathogenesis. Med (N Y). 2020.

39. Zheng ZQ, Wang SY, Xu ZS, Fu YZ, Wang YY. SARS-CoV-2 nucleocapsid protein impairs stress granule formation to promote viral replication. Cell Discov. 2021;7(1):38.

40. Cong Y, Ulasli M, Schepers H, Mauthe M, V’Kovski P, Kriegenburg F, et al. Nucleocapsid Protein Recruitment to Replication-Transcription Complexes Plays a Crucial Role in Coronaviral Life Cycle. J Virol. 2020;94(4).

41. Tugaeva KV, Hawkins DEDP, Smith JLR, Bayfield OW, Ker D-S, Sysoev AA, et al. The Mechanism of SARS-CoV-2 Nucleocapsid Protein Recognition by the Human 14-3-3 Proteins. Journal of Molecular Biology. 2021;433(8):166875.

42. Surjit M, Kumar R, Mishra RN, Reddy MK, Chow VT, Lal SK. The severe acute respiratory syndrome coronavirus nucleocapsid protein is phosphorylated and localizes in the cytoplasm by 14-3-3-mediated translocation. J Virol. 2005;79(17):11476–86.

43. Liu Y, Liu J, Plante KS, Plante JA, Xie X, Zhang X, et al. The N501Y spike substitution enhances SARS-CoV-2 transmission. bioRxiv. 2021.

44. Josset L, Menachery VD, Gralinski LE, Agnihothram S, Sova P, Carter VS, et al. Cell host response to infection with novel human coronavirus EMC predicts potential antivirals and important differences with SARS coronavirus. mBio. 2013;4(3):e00165–13.

45. Vanderheiden A, Edara VV, Floyd K, Kauffman RC, Mantus G, Anderson E, et al. Development of a Rapid Focus Reduction Neutralization Test Assay for Measuring SARS-CoV-2 Neutralizing Antibodies. Curr Protoc Immunol. 2020;131(1):e116.

46. Langmead B, Salzberg SL. Fast gapped-read alignment with Bowtie 2. Nat Methods. 2012;9(4):357–9.

47. Smith T, Heger A, Sudbery I. UMI-tools: modeling sequencing errors in Unique Molecular Identifiers to improve quantification accuracy. Genome Res. 2017;27(3):491–9.

