## supporting info for "Nucleocapsid mutations in SARS-CoV-2 augment replication and pathogenesis"

**Supporting Information**

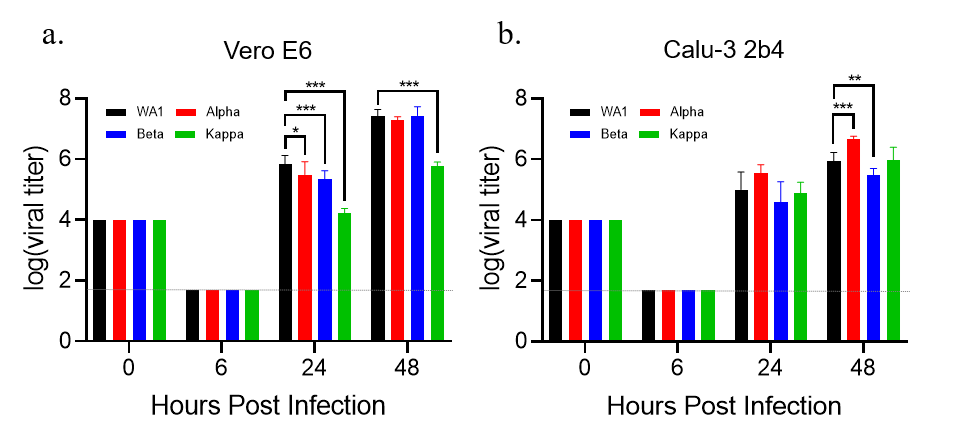

**S1 Fig. Replication of SARS-CoV-2 variants.** Viral titer from Vero E6 (**A**) or Calu-3 2b4 cells (**B**) inoculated with SARS-CoV-2 WA-1 (black) or the alpha (red), beta (blue) or kappa (green) variants at a MOI of 0.01. Graphed data represent the mean ± s.d. Statistical significance was determined by two-tailed student’s T-test with p≤0.05 (*), p≤0.01 (**), and p≤ 0.001 (***). Grey dotted lines are equal to LOD.

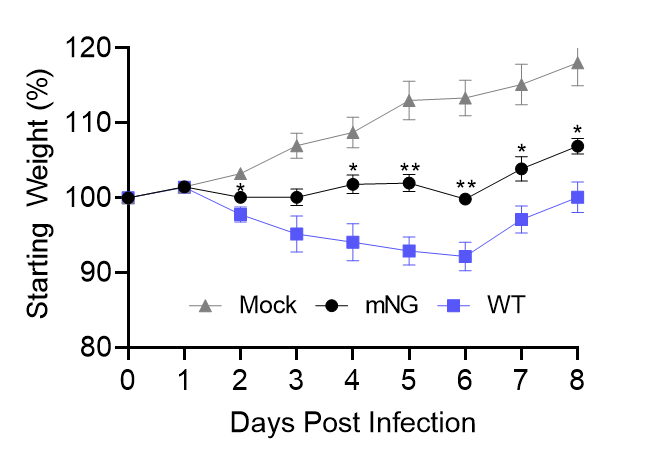

**S2 Fig. *In vivo a*ttenuation of the SARS-CoV-2 mNeonGreen reporter virus.** Three- to four-week-old Golden Syrian hamsters were intranasally inoculated with PBS alone (gray) or 10^4^ PFU of WA-1 SARS-CoV-2 (blue) or mNG SARS-CoV-2 (black). Graphed data represent the mean weight loss ± s.e.m (n≥5). Statistical significance between WT and mNG determined by two-tailed students T-test with p≤0.05 (*) and p≤0.01 (**).

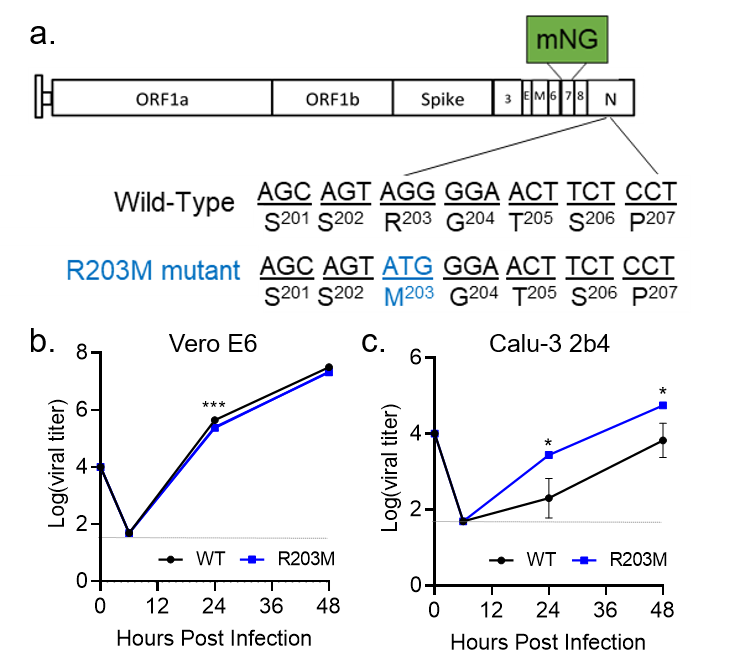

**S3 Fig. The R203M mutation enhances SARS-CoV-2 replication.** (**A**) Schematic of the SARS-CoV-2 genome, showing the creation of the R203M mutation and the replacement of ORF7 with the mNG reporter protein. (**B-C**) Viral titers from Vero E6 (B) or Calu-3 2b4 (C) infected with WT or R203M SARS-CoV-2 at an MOI of 0.01. Graphed data represent mean ± s.d. (n=3). Statistical significance was determined by two-tailed student’s T-test with p≤0.05 (*) and p≤ 0.001 (***). Grey dotted lines are equal to LOD.

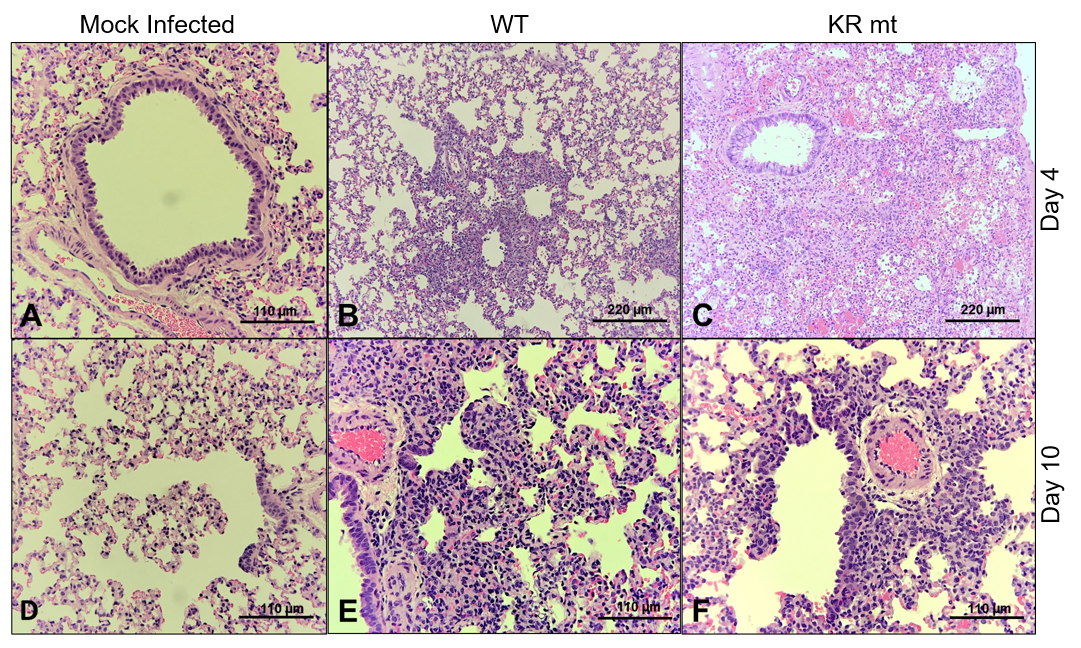

**S4 Fig. Lung histopathology in hamsters infected with WT and KR mt SARS-CoV-2.** Lung tissue was harvested, fixed, and 5 µm sections cut from mock, WT SARS-CoV-2, or KR mt-infected hamsters and stained with hematoxylin and eosin. (**A**) Normal bronchus, pulmonary artery, and alveoli in mock infection on day 4 (20X). (**B**) Bronchiolitis, peribronchiolitis, interstitial pneumonia, and edema surrounding branch of the pulmonary artery at day 4 in hamsters infected with WT virus (10X). (**C**) Severe bronchiolar cytopathic effect, interstitial pneumonia, cytopathic alveolar pneumocytes, alveoli containing mononuclear cells and red blood cells at day 4 in hamsters infected with KR mt. This lesion extended over numerous fields (10X). (**D**) Normal respiratory bronchiole, alveolar ducts, and alveolar sacs in mock infection on day 10 (20X). (**E**) Interstitial pneumonia adjacent to a bronchus at day 10 in hamsters infected with WT (20X). (**F**) Bronchiolar epithelial cytopathic effect, peribronchiolitis, focal interstitial pneumonia, branch of pulmonary artery with surrounding edema and mononuclear cell infiltration of endothelium at day 10 in a hamster infected with the KR mt (20X). Shown are representative images typical of data gathered from 5 animals from each group.

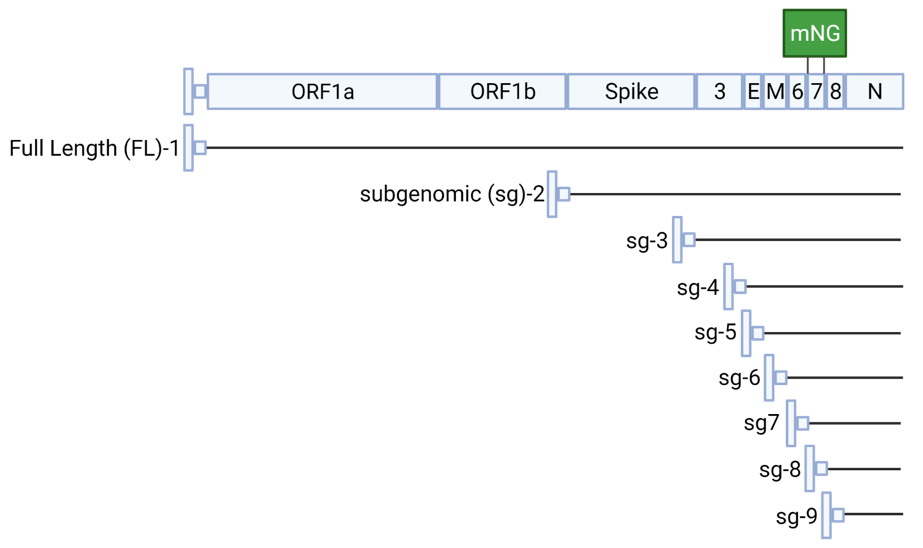

**S5 Fig. Schematic of SARS-CoV-2 RNAs.** Illustration of full length (FL) and subgenomic (sg) RNAs produced during SARS-CoV-2 infection.

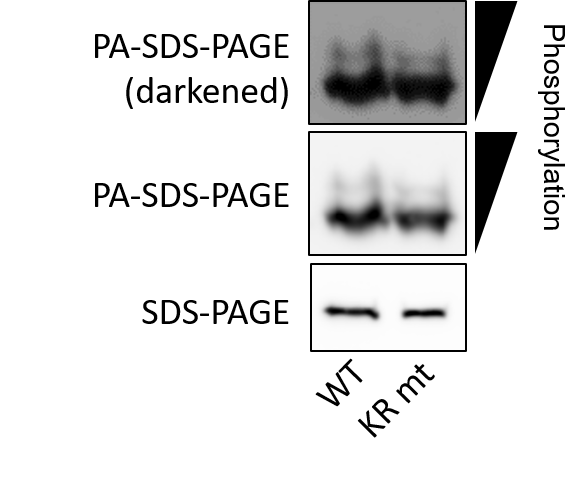
**S6 Fig.** **The KR mt has no effect on phosphorylation in virions**. Calu-3 2b4 cells were infected at an MOI of 0.01 with WT or KR mt SARS-CoV-2. Forty-eight hours post infection, viral supernatants were taken. Virions were then purified from supernatants by ultracentrifugation on a 20% sucrose cushion, inactivated, and N levels analyzed by both phospho-affinity and standard SDS-Page. Results are representative of two independent experiments.

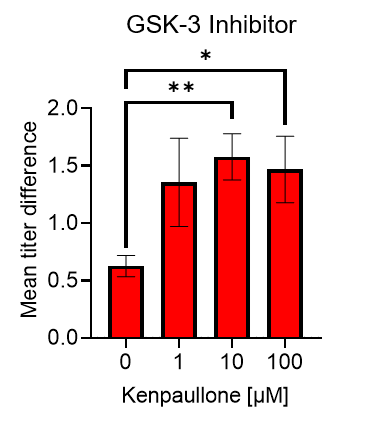

**S7 Fig. Mean titer differences between WT and KR mutant during GSK-3 inhibition**. Mean difference in viral titer between WT SARS-CoV-2 and the KR mutant when treated with kenpaullone at the indicated concentrations (n=4). Error bars are ± s.e.m. Significance by student T-test with p≤0.05 (*) and p≤0.01 (**).

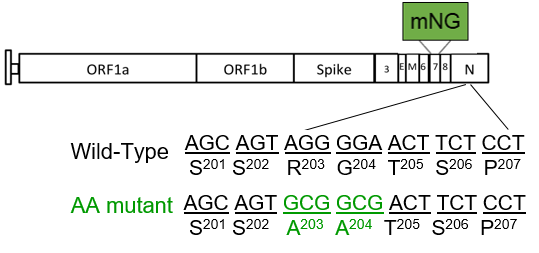

**S8 Fig. Schematic representation of the AA mt.** Schematic shows the creation of the AA mutation within the SARS-CoV-2 genome and the replacement of ORF7 with the mNeonGreen reporter.

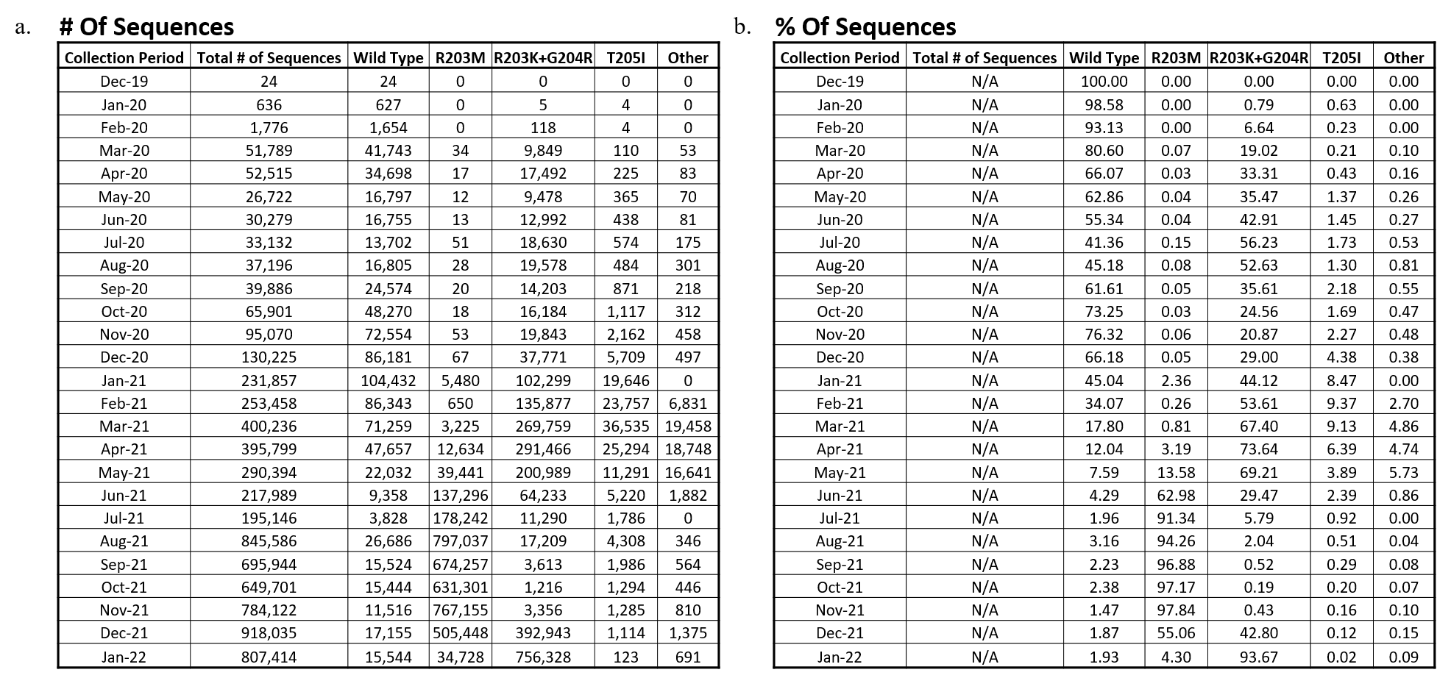

**S1 Table. Frequency of mutations in residues 203-205 in SARS-CoV-2 nucleocapsid.** (**A-B**) Frequency of WT, R203M, R203K+G204R, T205I, or all other genotypes binned by month of collection, represented as the raw totals (A) or as a percentage of total sequences in a given month (B).
